# Scaffolding protein GspB/OutB facilitates assembly of the *Dickeya dadantii* type 2 secretion system by anchoring the outer membrane secretin pore to the inner membrane and to the peptidoglycan cell wall

**DOI:** 10.1101/2021.08.06.455404

**Authors:** Shiheng Zhang, Shuang Gu, Piers Rycroft, Florence Ruaudel, Frederic Delolme, Xavier Robert, Lionel Ballut, Richard W. Pickersgill, Vladimir E. Shevchik

## Abstract

The phytopathogenic proteobacterium *Dickeya dadantii* secretes an array of plant cell wall degrading enzymes and other virulence factors via the type 2 secretion system (T2SS). T2SSs are widespread among important plant, animal and human bacterial pathogens. This multi-protein complex spans the double membrane cell envelope and secretes fully folded proteins through a large outer membrane pore formed by 15 subunits of the secretin GspD. Secretins are also found in the type 3 secretion system and the type 4 pili. Usually, specialized lipoproteins termed as pilotins assist the targeting and assembly of secretins into the outer membrane. Here, we show that in *Dickeya*, the pilotin acts in concert with the scaffolding protein GspB. Deletion of *gspB* profoundly impacts secretin assembly, pectinase secretion, and virulence. Structural studies reveal that GspB possesses a conserved periplasmic Homology Region domain that interacts directly with the N-terminal secretin domain. Site-specific photo cross-linking unravels molecular details of the GspB-GspD complex *in vivo*. We show that GspB facilitates outer membrane targeting and assembly of the secretin pores and anchors them to the inner membrane while the C-terminal extension of GspB scaffolds the secretin channel in the peptidoglycan cell wall. Phylogenetic analysis shows that in other bacteria, GspB homologs vary in length and domain composition and act in concert with either a cognate ATPase GspA or a pilotin GspS.

**Importance:** Gram-negative bacteria have two cell membranes sandwiching a peptidoglycan net that form together a robust protective cell envelope. To translocate effector proteins across this multilayer envelope, bacteria have evolved several specialized secretion systems. In the type 2 secretion system and some other bacterial machineries, secretins form large multimeric pores that allow transport of effector proteins or filaments across the outer membrane. The secretins are essential for nutrient acquisition and pathogenicity and constitute a target for development of new antibacterials. Targeting of secretin subunits into the outer membrane is often facilitated by a special class of lipoproteins called pilotins. Here, we show that in *Dickeya* and some other bacteria, the scaffolding protein GspB acts in concert with pilotin, facilitating the assembly of the secretin pore and its anchoring to both the inner membrane and the bacterial cell wall. GspB homologs of varied domain composition are present in many other T2SSs.

## Introduction

Gram-negative bacteria are surrounded by two membranes, the inner membrane (IM) and the outer membrane (OM) that together delimit a thin (∼30 nm) periplasmic space containing the peptidoglycan (PG) layer (1, 2). To exchange proteins, nucleic acids and sugars with the external medium, these bacteria have evolved an array of specialized transport systems (3-7). The type 2 secretion system (T2SS) is widespread in proteobacteria and secretes folded proteins that play a pivotal role in colonization of different niches, survival, competition and pathogenicity (8-12). The plant pathogenic γ-proteobacterium *D. dadantii* uses the T2SS, called Out, to secrete pectinases in the infected plant tissues causing soft rot disease in numerous plants and root-vegetables (13, 14). The T2SS is embedded in both the IM and the OM and spans the whole cell envelope of the bacteria. It is composed of 12 core components, generically called GspC to GspM and GspO (OutC to OutO, in *D. dadantii*) as well as some additional components, GspA, B, N and S, that are present in certain T2SSs (8-10, 15, 16).

The secretin is an essential T2SS component that forms gated channels in the OM, through which the effector proteins are translocated in the medium or at the cell surface. The secretins are also shared by the type 3 secretion system (T3SS), type 4 pili (T4P) as well as by the competence and the filamentous phage assembly systems (16-19). Secretin homologs have also been identified in mitochondria of some eukaryotes (20). Recent cryo-EM structures have revealed important molecular details of the assembly of the MDa-sized secretin channels consisting of 12 to 16 subunits, with a clear predominance of 15-fold symmetry in more recent, near-atomic resolution structures of the T2SS and T3SS secretins (15, 21-28). Only a small apical segment of the secretin channel consisting of an amphipathic helical loop and a ‘β-lip’ is embedded into the OM, while the main portion, which is composed of the core secretin domain together with the N-terminal domains, N0 to N3, are located in the periplasm. The conserved C-domains of 15 secretin subunits build up together a double β-barrel channel composed of 60 internal and 60 external β-strands. The N1 to N3 domains also termed as ring-building motifs, adopt mixed α/β fold and form together a cylinder like structure protruding through the periplasm. In the T2SS secretins, the N-terminal N0 domain forms the first gate at the entry of the secretin channel that interacts with the IM portion of the secretion machinery and controls the recruitment of substrates (29-33). This portion of the secretin channel is not seen in the majority of reported structures, or appeares as smeared density consistent with its flexibility (21, 23, 34). However, in the recent high-resolution cryo-EM structure of *Klebsiella pneumoniae* PulD, the N0-N1 domains have been resolved as a tightly packed ring, stabilized by interaction with the inner membrane PulC component (27). The molecular details of this interaction remain elusive, but this study shows the relevance of attachment of the secretin channel to the periplasmic portion of the IM components.

Usually, a small lipoprotein called pilotin guides the cognate secretin subunits through the periplasm to the OM and facilitates their assembly (16, 35). Two groups of sequence and structure-dissimilar pilotins have been identified in various T2SSs. The paradigm pilotins of the OutS/PulS family from *Dickeya, Klebsiella* and enterohemorrhagic *E. coli* bind a short C-terminal region of secretin, termed as S-domain, and pilot the secretin subunit via the Lol system to the inner leaflet of the outer membrane (36-39). Upon binding of pilotin, the disordered S-domain folds into α-helix (40-42). In the assembled secretin channel, the bound pilotins stabilize the external β-barrel by stapling tandemly arranged secretin subunits (15, 43). The T2SS pilotins of the AspS/ExeS family that were identified in *Vibrio, Aeromonas*, enterotoxigenic and enteropathogenic *E. coli* seem to fulfil exactly the same functions than OutS/PulS while adopting profoundly different fold (15, 43-45). Targeting and assembly of secretins from the T3SS and T4P systems are also facilitated by specific piloting lipoproteins of different structures and modes of action. For instance, the T3SS pilotins MxiM and ExsB are structurally unrelated neither to one another, nor to PilF/PilW family pilotins from the T4P system, or to the T2SS pilotins (46-49).

Beside the pilotins, some other assistance proteins are also involved in the assembly of the secretin channels through the bacterial cell wall. The main portion of the secretin channel is located in the periplasm (18 to 20 nm) and consequently, it crosses through and could interact with the PG mesh (27, 34). Indeed, the distance between the OM and the PG layer is 12 to 25 nm and is controlled by the size of the major OM lipoprotein Lpp, or Braun’s lipoprotein, that attaches covalently the PG to the OM (1, 50). In the high-resolution cryo EM tomography of the *Salmonella typhimurium* T3SS and the *Myxococcus xanthus* T4P, the PG layer was visualized around the N-terminal portion of the respective secretin channels (25, 51). In the *M. xanthus* T4P, the N0 domain of the secretin PilQ is preceded by a triplet of specialized PG-binding AMIN domains (25, 52). In addition, another T4P component, TsaP, carries a LysM-like PG-binding domain; together they provide anchoring of the secretin channel to the PG (53). The PG-binding modules have been identified in the other trans-envelope machineries of Gram-negative bacteria illustrating that anchoring to the PG is essential for their assembly and function (54-56).

In the T2SS, only LspD secretin from *Legionella* is known to possess a specialized PG-binding SPOR domain, located at the N-terminus just prior to the N0 domain (34). Another known example of PG-binding T2SS component is the multimodular ATPase GspA that possesses a periplasmic PG-binding domain of Pfam family PF01471 (57). GspA has been identified in *Vibrio, Aeromonas* and some other bacteria, where it acts together with GspB and is thought to be implicated in the insertion of the secretin into or through the PG mesh (58, 59). GspB is also present in some other *Enterobacteriaceae*, such as *Dickeya, Klebsiella* and *Pectobacterium*, but in this case, without a GspA counterpart (60). Previous studies have suggested that GspB could interact with the secretin but its precise role has remained unclear and conflicting (60-63).

In this study, we explore the structure and function of OutB/GspB from plant pathogen *D. dadantii*, we reveal that the GspB periplasmic domain is structurally similar to the homology region (HR) of GspC and depict the molecular details of its interaction with the N0 domain of the secretin. We demonstrate that OutB/GspB guides the secretin to the outer membrane and anchors the secretin to the inner membrane and to the cell wall peptidoglycan.

## Results

### OutB is required for type 2 secretion and essential for the full virulence of *D. dadantii*

Collective action of several pectinases secreted by *D. dadantii* Out T2SS causes soft-rot maceration of infected plants, while mutant bacteria lacking functional Out system are fully or partially avirulent (14, 64). Therefore, to examine the functional relevance of OutB, we assessed its importance in the context of plant infection. Pathogenicity assays with chicory leaves clearly show that the *D. dadantii outB* deletion mutant was barely virulent in comparison with the wild-type strain. A very small rotted area formed by *outB* mutant 24 hours post infection did not progress further and was surrounded by dehydrated necrotic plant tissues that form a barrier for bacterial proliferation (Fig. 1A). *D. dadantii outD* mutant, which lacks the secretin pore and was used as a negative control, provoked very similar symptoms, with an even smaller infection zone. Since soft-rot symptoms result from the action of pectinases secreted by the Out system, these data indicate that *outB* and *outD* mutants were both unable to efficiently secrete pectinases into the plant tissue. Consistent with this, in plate secretion assays that show pectin and cellulose degradation by the Out-secreted enzymes, the *outB* mutant generated no or very small halos, similar to the *outD* mutant (Fig. 1B). These data show a loss or malfunction of the Out T2SS in the *outB* mutant that causes a striking reduction of bacterial virulence *in planta*.

**Figure 1.**
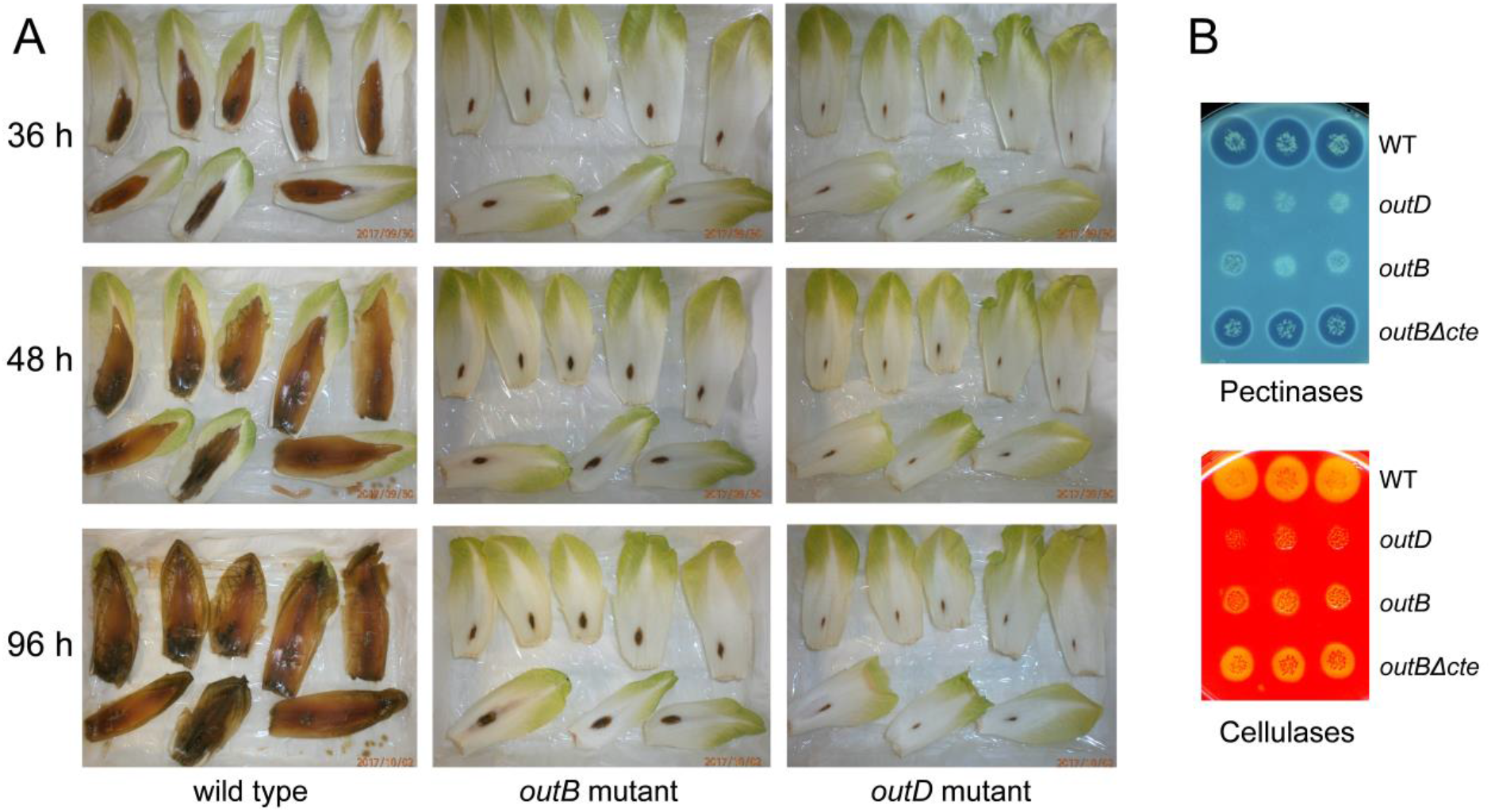
OutB is required for the type 2 secretion and full virulence of *D. dadantii*. **A**. Pathogenicity tests with chicory leaves. Chicory leaves were inoculated with 10^6^ cells of *D. dadantii* wt, *outB* or *outD* strains and incubated at 28°C for the indicated time. **B**. Pectinase and cellulase secretion assays. *D. dadantii* strains carrying the indicated mutations were grown for 14 h at 30°C on plates containing polygalacturonate or carboxymethylcellulose and flooded with cupper acetate or Congo Red (upper and lower panels, respectively). Size of halo reflects the level of pectinase and cellulase secretion, respectively.

### OutB is attached to both the inner and the outer membranes

To examine if the absence of OutB compromises the integrity of the T2SS, the quantity of various T2SS components was assessed by immunoblotting (Fig. S1). The abundance of the inner membrane assembly platform component OutC and the major pseudopilin OutG did not vary significantly, while the quantity of the secretin OutD decreased in the *D. dadantii outB* mutant. Conversely, the amount of OutB was obviously lower in *D. dadantii outD* strain (Fig. S1). These data indicate that OutB is required for secretin channel biogenesis or stability. OutD is localized in the OM while OutB is predicted to reside in the IM, due to its hydrophobic N-terminal segment (Fig. 2A). To test how these proteins could interact within the bacterial envelope, we fractionated membrane vesicles from *D. dadantii* wild type cells on a sucrose gradient (Fig. 3). No full-length OutD but an abundant OutD cross-reacting band of ∼35 kDa was detected in the OM fractions (Fig. 3A) suggesting OutD degradation during the 60 h centrifugation (probably by the *D. dadantii* metalloproteases (65) since EDTA could not be used in the course of membrane separation). OutB co-localized with both membranes, in contrast to the *bona fide* IM protein TolA and the OM components, OmpA porin and lipopolysaccharide, LPS. This suggests that a fraction of OutB is attached to the OM or to an OM-associated component, for example, the secretin or the peptidoglycan. The treatment of cell extracts with lysozyme, to release the OM vesicles from the PG mesh, improved the separation of the OM components, LPS and OmpA, which moved to the higher density fractions, but OutB remained split between the two membranes (Fig. 3B). However, when the membranes of a *ΔoutD* strain were separated, the effect of the PG on OutB location became clearly visible (Fig. 3C and D). Here, and only after cell-wall cleavage by lysozyme, OutB was detected uniquely in the IM fractions. These data suggest that OutB attaches to both the PG and OutD. We further examined these two possibilities.

**Figure 2.**
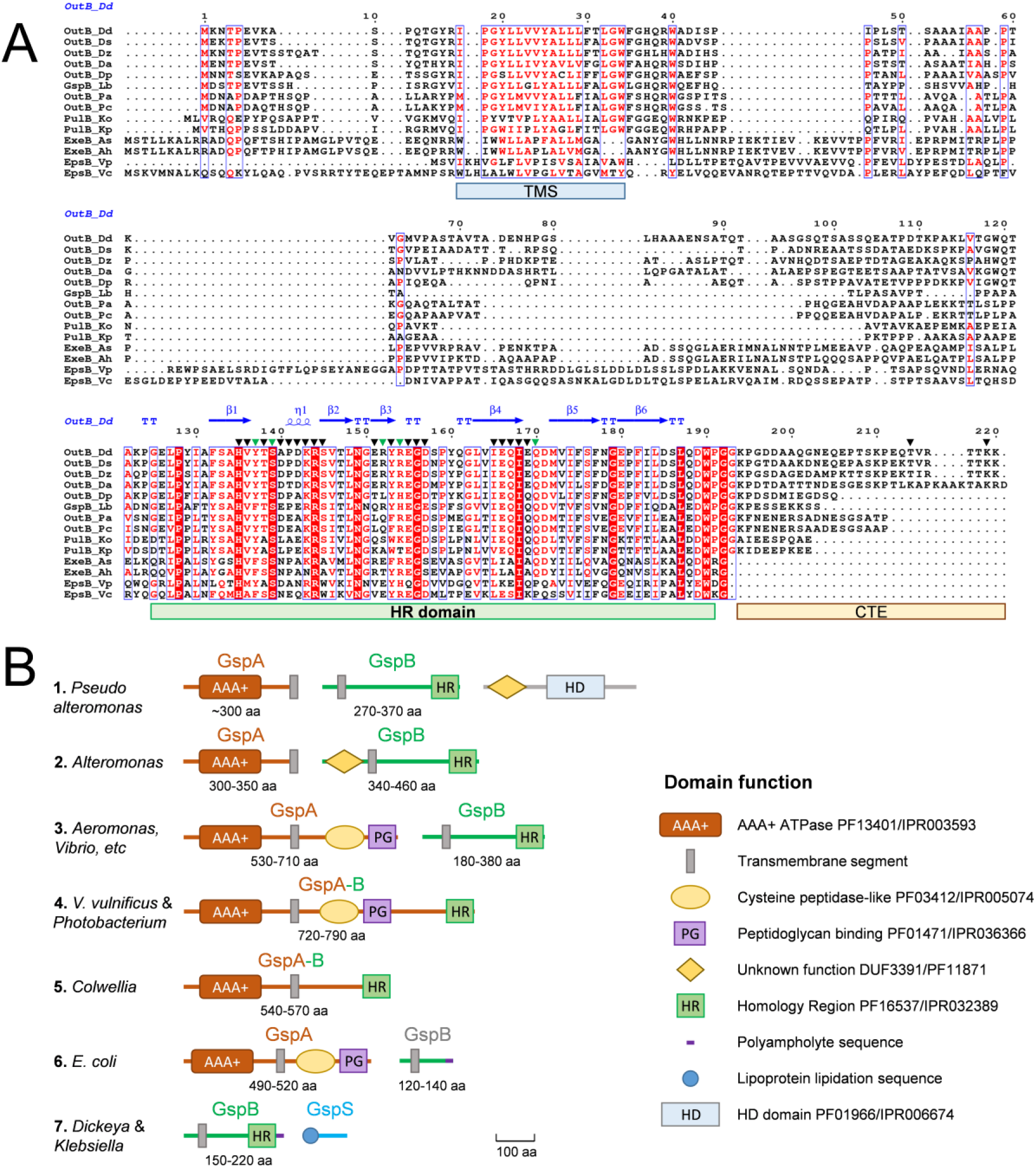
GspB organization. **A**. Alignment of representative GspB sequences. Multiple sequence alignment was performed with Clustal Omega (ebi.ac.uk) and ESPript server (107). The secondary structure elements are shown for the OutB HR domain (PDB entry 4WFW). The residue numbering is this for *D. dadantii* OutB. The groups of identical and similar residues are highlighted in red or are in red, respectively. The residues substituted with *p*BPA and used in photo-cross-linking are shown with triangles; with green triangles are the residues generating an abundant complex with OutD. Positions of the transmembrane segment (TMS), the homology region (HR) and the C-terminal extension (CTE) are indicated with coloured bars. Accession numbers of the used protein sequences are listed in Table S6. **B**. Typical gene and domain organizations of GspB and related GspA and GspS proteins. For each GspB representative shown in Fig. S14, gene synteny and domain organization were analyzed with NCBI (www.ncbi.nlm.nih.gov), InterPro (www.ebi.ac.uk/interpro) and Pfam (pfam.xfam.org) data bases and summarized into seven architypes, named according to the most abundant or most studied representative bacteria. The protein size range (in amino acids) is indicated for each group. Of note is that more than 80% of analyzed GspB proteins belong to the archetype #3.

**Figure 3.**
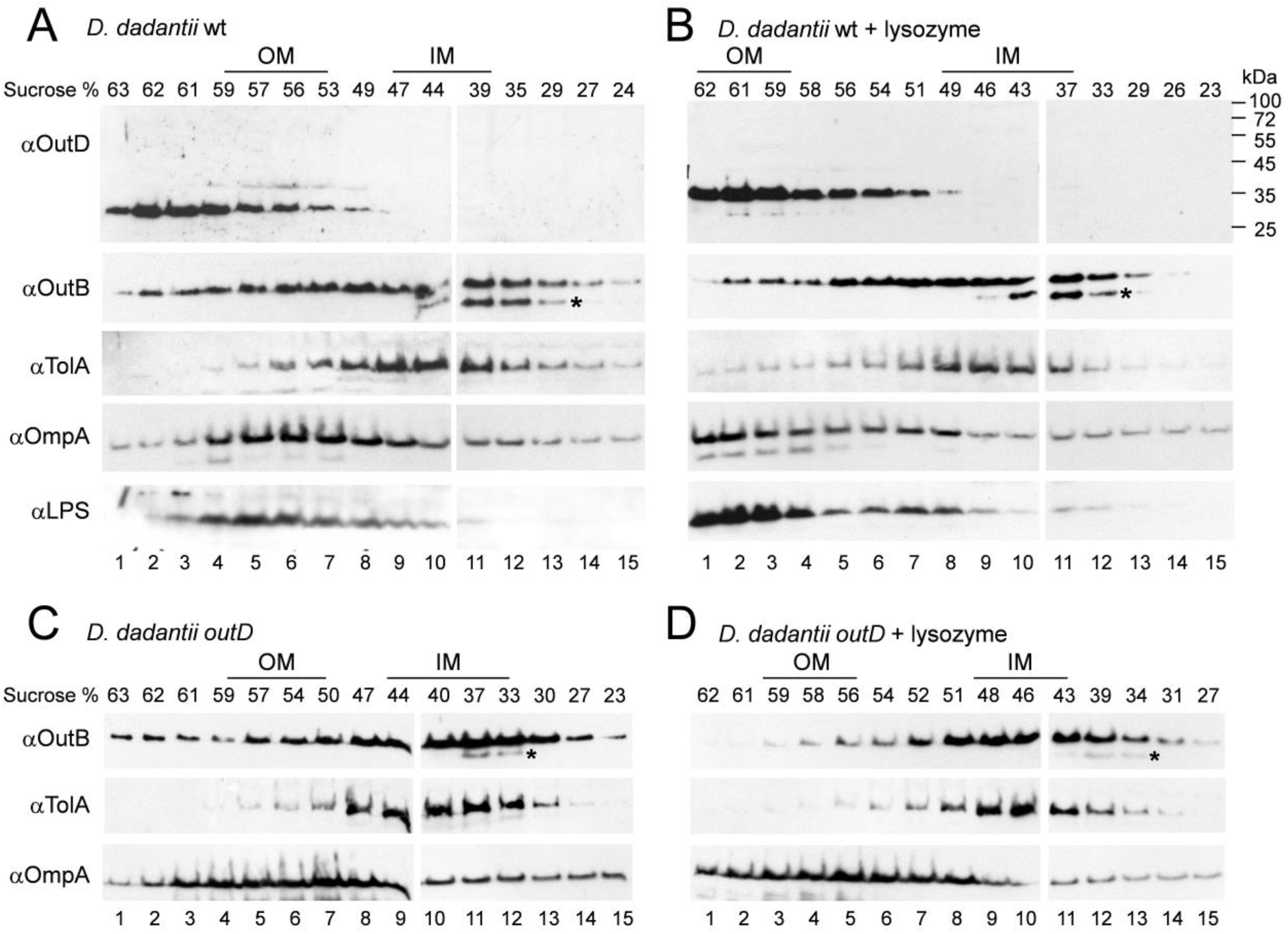
OutB is associated with both the inner and the outer membranes. *D. dadantii* A5652 wt (**A** and **B**) and A5653 *outD* (**C** and **D**) cells were grown on pectin containing plates, broken in French press and membranes were separated on sucrose gradient and analyzed by immunoblotting with the indicated antibodies. In **B** and **D**, the broken cell extracts were treated with lysozyme prior loading onto the gradient. The position of the inner and the outer membrane fractions are indicated according to the positions of LPS and OmpA (OM) as well as TolA (IM). A degradative product of OutB is indicated with an asterisk.

### OutB suppresses PspA induction caused by mislocated secretin

Inefficient targeting of secretins to the OM results in their spontaneous insertion into the IM that causes ion leakage from the cytoplasm. This provokes induction of <u>p</u>hage <u>s</u>hock <u>p</u>rotein (Psp) stress response system that acts to restore membrane integrity (37, 66-70). Consequently, an increased level of PspA is an indicator of inefficient targeting of secretins. In *D. dadantii*, PspA is induced in the absence of the pilotin OutS, which promotes OutD targeting to the OM (40, 68).

To examine if OutB could also assist the OutD targeting and affect the PspA response, *outD* was expressed in *E. coli* either alone or with *outB* and/or *outS* from constructs designated as D, DB, DBS and DS (Fig. S2). As expected, OutS caused a significant increase of the OutD level and reduced that of PspA (Fig. 4A, compare DS with D). In contrast, OutB did not consistently affect the amount of OutD but notably reduced the PspA response, and that, in a manner independent on OutS (Fig. 4A, compare DB and DBS with D and DS). These data suggest that OutB could improve OutD targeting to the OM. Consistent with this possibility, sucrose gradient analysis showed a lower proportion of OutD in the IM fractions of the DBS sample in comparison with DS (Fig. 4B). In addition, the total amount of OutD was visibly lower in the DS gradient, suggesting that in the absence of OutB, OutD is more prone to degradation during the gradient separation (Fig. 4B). Therefore, OutB protects OutD from degradation and facilitates its transport to the OM.

**Figure 4.**
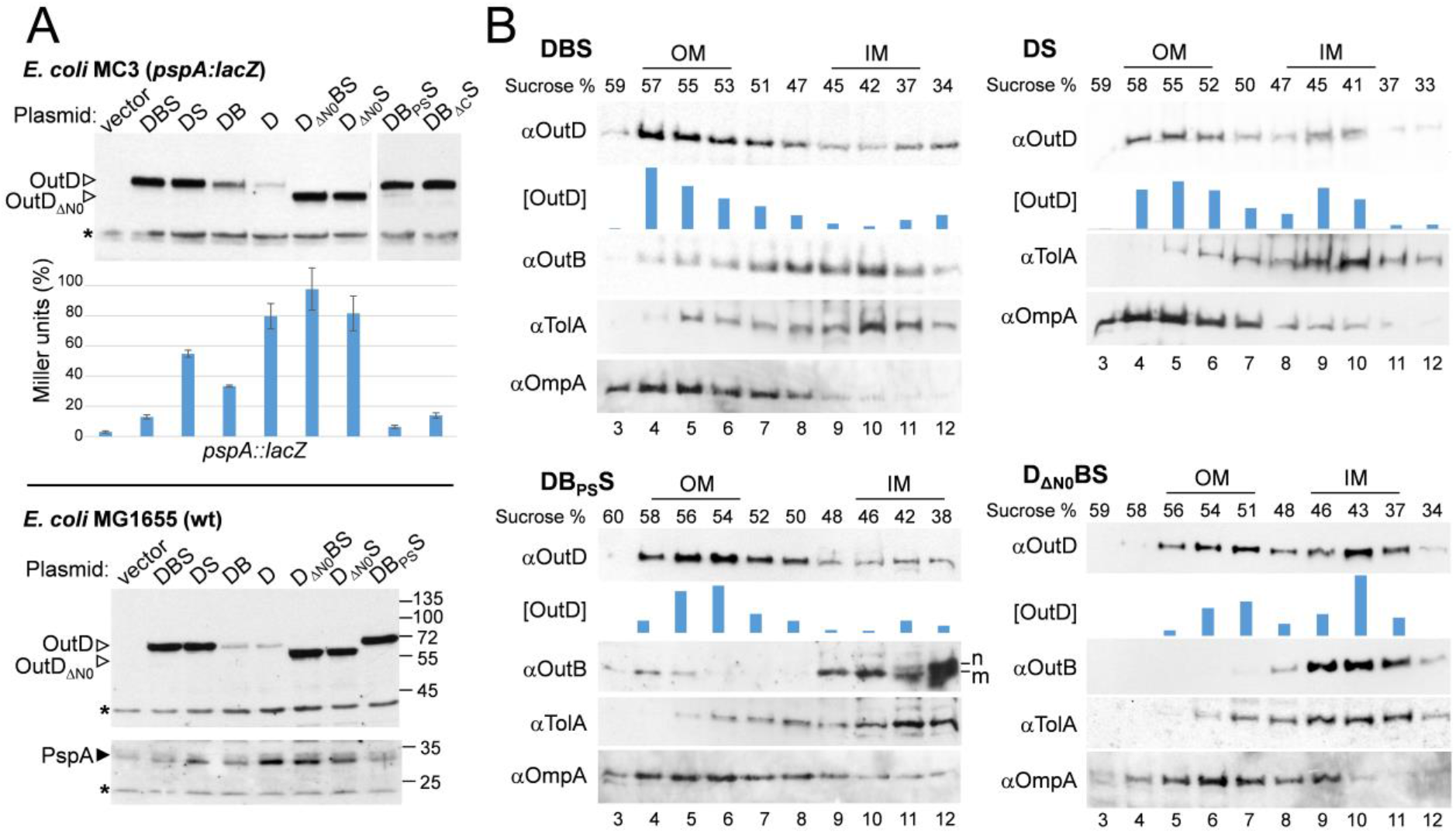
OutB suppresses PspA response caused by OutD and assists OutD targeting in the OM. **A**. *E. coli* MC3 (upper panels) and MG1655 (lower panels) carrying pREP4 (*lacI*^*Q*^) and pGEMT plasmid with the indicated combinations of *outD, outB* and *outS* (D, B and S, respectively) were grown at 28°C for 14 h. The level of OutD and PspA was probed by immunoblotting or by β-galactosidase activity measurement, for *pspA::lacZ*. The β-galactosidase values are the mean of three cultures; one of which was also probed by immunoblotting with anti-OutD. Asterisk indicates a nonspecific cross-reacting protein, used as a loading control. **B**. *E. coli* MC3 cells grown as in ***A*** were broken in French press and membranes were separated on sucrose gradient and analyzed by immunoblotting with the indicated antibodies. The positions of IM and OM are indicated according to the location of TolA and OmpA, respectively. The quantity of OutD in the fractions, [D], was estimated with Image Lab software (Bio-Rad). The exposition time of DS panel with OutD antibodies was 10 min versus 3 min with DBS panel, consistent with a lower OutD content. Non-matured and matured versions of OutB_PS_ are indicated with ‘n’ and ‘m’.

### The periplasmic OutB region decreases the PspA response caused by OutD mislocation

OutB is composed of a short cytoplasmic region, a transmembrane helix followed in the periplasm by a linker region of variable length, and a conserved periplasmic domain, ended with a 28-residue long C-terminal extension (CTE) (Fig. 2A). We examined which part of OutB is critical for reduction of the PspA response. When the CTE was deleted, the resulting OutB_*Δ*CTE_ reduced a PspA level similar to that with the full-length OutB, indicating that this part of OutB is not essential in this context (Fig. 4A, compare DB_*Δ*C_S and DBS). In contrast, when the transmembrane segment of OutB was replaced with the cleavable N-terminal signal peptide, the resulting OutB_PS_ reduced the PspA response even slightly better than did OutB (Fig. 4A, compare DB_PS_S and DBS). In the sucrose gradient of DB_PS_S, OutD co-migrated mainly with the OM fractions similarly to that in DBS (Fig. 4B). Notably, a small proportion of the matured, signal peptide-less OutB_PS_ co-migrated with the high-density OM indicating that the periplasmic, membrane segment-free OutB_PS_ binds OutD strongly enough to move together through the gradient.

### OutD N0 domain is required for secretin targeting and gating the secretin channel

In the reverse experiments, we examined which part of OutD is involved in the chaperoning by OutB. Expression of OutD_*Δ*N0_ lacking the N0 domain (missing 91 residues of matured OutD, from F7 to I96) elicited high PspA response even in the presence of OutB (Fig. 4A). Membrane separation shows that in contrast to the wild type OutD, more than one-half of OutD_*Δ*N0_ co-migrated with the IM fractions, (Fig. 4B, compare D_*Δ*N0_BS to DBS). In addition, in the D_*Δ*N0_BS gradient, OutB was strictly confined to a few IM fractions while in DBS (OutD_WT_), a significant proportion of OutB co-migrated with the intermediate and the OM fractions. These data show that in the absence of the N0 domain, OutB is no longer able to bind OutD and assist its correct targeting.

### OutB HR domain interacts with OutD N0 domain

A possible direct interaction between OutB and OutD was first investigated by GST-pull down assay. GST-tagged C-terminal portion of OutB consisting of HR and CTE (residues P112 to K220) bound very efficiently the OutD derivatives comprising N0-N1-N2 domains and to a lone N0 domain but not to the N1-N2 derivative (Fig. S3). Deletion of the 28-residue CTE did not affect the interaction, narrowing the interacting domains down to the HR domain of OutB (residues P112 to G192) and N0 domain of OutD (residues A1 to S85) (Fig. S3).

### NMR characterization of OutB HR/OutD N0 complex

NMR spectroscopy was further used to explore the interaction of the OutB HR and OutD N0 domains. Initial HSQC experiments using ^15^ N-labelled HR domain of OutB (residues 112 to 220) and two OutD derivatives, OutD-N0 (residues 1-85) and OutD-N0,N1,N2 (residues 1-258), showed that both OutD derivatives gave the same pattern of peak shifts confirming that it is the N0 domain of the secretin that interacts with the HR domain of OutB (Fig. S4). Partial assignment of the ^1^ H-^15^ N HSQC spectra for OutB HR domain showed that several residues from different zones on the surface of the HR domain are affected by the interaction with the OutD N0 domain (Fig. S5A). In reciprocal experiments, residues of the OutD N0 domain involved in the interaction surface could not be readily identified from peak shifts, because when the OutB HR domain was titrated into ^15^ N-labelled N0 domain there were widespread chemical shift changes (Fig. S5B). Such peak shifts are occasionally seen when the electron distribution is perturbed on forming a complex. Determination of the interdomain interface in the OutB HR/OutD N0 complex by NMR experiments was therefore not straightforward.

### Crystal structure of the periplasmic domain of OutB

The high resolution structures of the N0 GspDs domain have been reported for *E. coli* and *P. aeruginosa* (31, 71). To define the molecular details of the OutB HR/OutD N0 interaction, we sought to solve the structure of the OutB HR domain. Three constructs were used in crystallization trials: OutB^112-220^, OutB^112-202^ and OutB^112-192^ of which OutB^112-202^ gave crystals used to determine the structure at 2.05 Å resolution (Table S1). The structure of the OutB HR domain was solved using the enterotoxigenic *E. coli* HR GspC domain as a search model in molecular replacement (17% of sequence identity) (31). Residues 115-198 of OutB are clearly defined in the electron density map. The HR domain of GspB consists of two three-stranded antiparallel β-sheets; the β-strands sequentially form the up-down-up β-sheets, so the first sheet comprises strands 1, 2 and 3 and the second, 4, 5 and 6 (Fig. 5A). The two sheets are at approximately 70° to each other, such as the structure forms a β-sandwich with a hydrophobic core. The 3_10_-helix in the loop between β1 and β2 contains a highly conserved R144, which forms a salt bridge to E155, also highly conserved (Fig. 2A and S6A). Perhaps the most striking feature of the structure is the quantity of irregular polypeptide, between the short β-strands, especially but not limited to the β3/ β4 loop, and at the amino- and carboxy-ends of the β-sandwich (Fig. 5A). DSSP algorithm (72) reveals 58% of the residues do not fall into a recognized secondary structure category.

**Figure 5.**
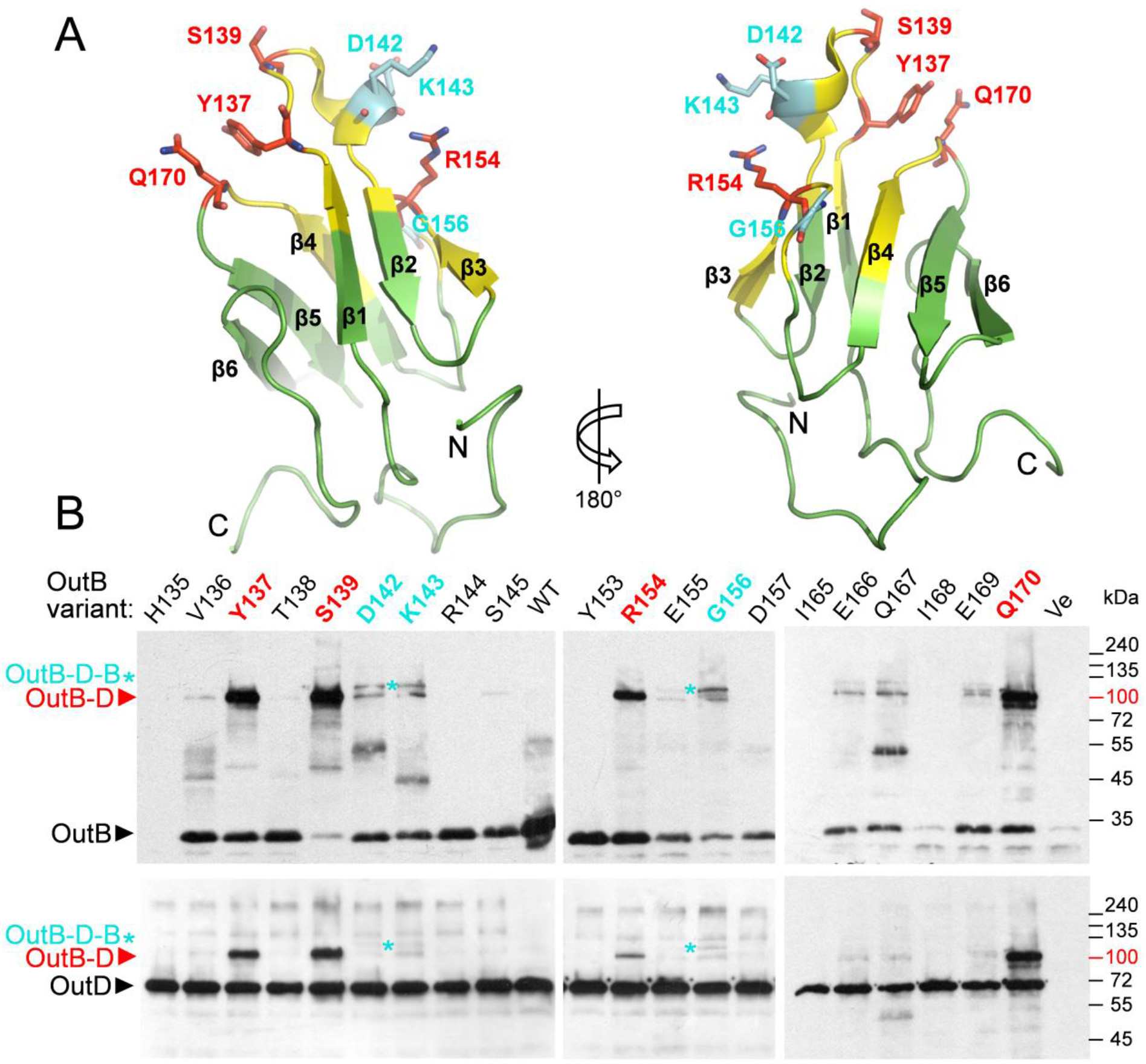
Photo-crosslinking maps OutB sites interacting with OutD. **A**. OutB HR structure is formed by two three-stranded anti-parallel β-sheets coming together to form a β-sandwich. The residues generating abundant OutB-D complexes are in red, those generating putative OutB-D-B complex are in cyan and other residues probed by photo-crosslinking are in yellow. **B**. *In vivo* site specific photo-crosslinking. OutB *p*BPA substitutions (indicated at the top) were expressed from DBS plasmid in *E. coli* MG1655/pREP4/pSupBPA. Cells were irradiated by UV (365 nm) for 3 min and analyzed by immunoblotting with anti-OutB and OutD antibodies, upper and lower panel, respectively. Only UV-irradiated cells extracts are shown. An equivalent amount of cells was loaded into each well. OutB, OutD and their complexes are indicated by an arrow or by an asterisk.

A search for structural homologs using the DALI webserver (73) returned the HR GspC domain from enterotoxigenic *E. coli* used as the molecular replacement search model as the most similar structure (PDB entry 3OSS, Z-score 7.8, r.m.s.d. 1.7Å for 58 equivalent CA atoms) (31). The other related architectures are PilP pilot protein from the T4P systems of *Neisseria meningitidis* (PDB entry 2IVW), *P. aeruginosa* (PDB entry 2LC4) and *D. dadantii* OutC HR (PDB entry 2LNV) (74-76) (Fig. S6B).

### *In vivo* mapping of OutB site interacting with OutD

To characterize the architecture of the OutB-OutD complex *in vivo*, site-specific photo cross-linking was employed. In this approach, a photo-reactive amino acid *para*-benzoyl-phenylalanine (*p*BPA) is incorporated *in vivo* in place of the residue of interest (77, 78). Upon a short UV exposure of cells, *p*BPA can cross-link to any of the carbon-hydrogen bonds within a distance of 3 Å. In this way, we introduced *p*BPA in place of 24 residues of the OutB HR domain and assessed their cross-linked patterns in *D. dadantii* and *E. coli* cells. An abundant adduct of 110 kDa, cross-reacting with OutB and OutD antibodies and compatible with an OutB-OutD complex, was generated by OutB carrying *p*BPA in place of S139, Y137, Q170 and R154 (ranked in a descending order of abundance) (Fig. 5B and S7). Remarkably, OutB_S139*p*BPA generated a near-quantitative cross-linking to OutD, indicating an important OutB-OutD contact. When OutD*Δ*N0 lacking the N0 domain was used in place of full-length OutD, no OutB-D complex was generated indicating that OutB HR interacts directly with OutD N0 (Fig. S8).

Notably, the substitutions located at or near the β1 strand of OutB HR generated cross-linking patterns with even/odd alternation, typical for a β-strand addition (79). For instance, an OutB-D complex was generated with *p*BPA substitutions of Y137 and S139 but not with V136, T138 or A140 (Fig. 5B and S7). These data suggest that the β1 strand of OutB HR interacts with a β-strand from OutD N0. Since Q170 (β4-β5 turn) is spatially proximal to Y137 and S139 (β1-strand), these three residues seem to constitute the main OutD-interacting zone of OutB. Another residue generating an abundant OutB-OutD complex, R154, is located rather far from this patch of highly reactive residues (Fig. 5A) suggesting formation of an extended HR/N0 inter-domain interface or even two interfaces. Interestingly, some OutB_*p*BPA variants, e.g. D142, K143 and G156, generated two OutB and OutD cross-reactive adducts, one OutB-OutD and another one, of higher mass, compatible with OutB-OutD-OutB complex (Fig. 5B). The latter, putative ternary complex could result from a simultaneous cross-linking of one N0 domain to two HR domains.

### Mapping of OutD site interacting with OutB

To identify the OutD residues interacting with OutB HR, photo cross-linked complexes generated by OutB_*p*BPA variants S139, R154 and Q170 were purified by Strep-Tactin chromatography, separated by SDS-PAGE, followed by in-gel digestion with chymotrypsin and trypsin. The peptides were then analysed in a high resolution LC-MS/MS Q Exactive HF Mass Spectrometer (Thermo Fisher Scientific) (Fig. 6A and S9). LC-MS/MS data were submitted to StavroX software tools (80), which allowed an accurate identification of OutB_S139*p*BPA-OutD cross-linked peptides. Precisely, at a 1% False Discovery Rate (FDR), several best spectra with a score above the score cut-off of 48 correspond to the OutD peptide S^41^ YDMMNEGQY^50^, cross-linked to the OutB peptide YTS^139^ pBPAAPDKR^144^ (Fig. 6A and S10). The OutD residue, being cross-linked to *p*BPA, could be assigned from S^41^ to M^45^ with prevalence for D^43^ and M^44^ (Table S2).

**Figure 6.**
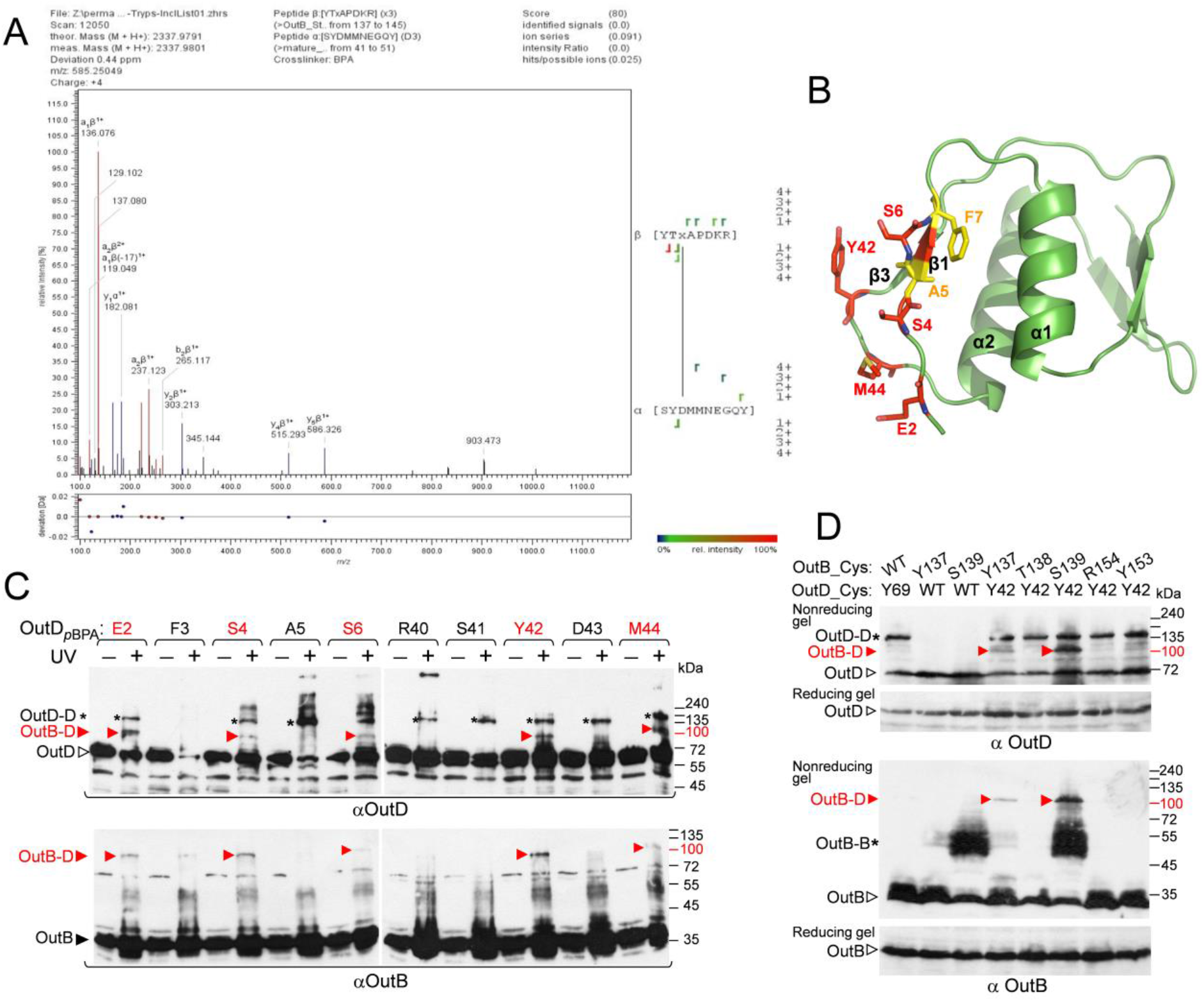
Mapping of the OutD site interacting with OutB. **A**. LC MS/MS analysis shows that OutB_S139*p*BPA interacts with SYDMMNEGQY peptide of OutD. OutB_S139*p*BPA-OutD complex was purified by Strep-Tactin chromatography followed by SDS-PAGE and in-gel digestion (Fig S9). The peptides were subjected to a high-resolution LC MS/MS analysis and MS spectra were assigned with StavroX. Shown is a representative MS/MS spectrum with the assigned OutD (α) and OutB (β) peptides and crosslinking site, where ‘x’ indicates the *p*BPA residue. The StavroX validation Score of 80 is above the score cut-off of 48 (Fig S10). **B**. OutD N0 residues probed by photo cross-linking: these generating OutB-D and OutD-D complexes are in red and in yellow, respectively. **C**. *In vivo* photo cross-liking maps the OutD residues interacting with OutB. OutD_*p*BPA substitutions (indicated at the top) were expressed from a DBS plasmid in *E. coli* MG1655/pREP4/pSupBPA. Cells were irradiated by UV and analysed by immunoblotting with OutB and OutD antibodies. **D**. Disulfide bonding analysis maps an OutB-OutD interacting site. *E. coli* MG1655/pREP4 carrying a DBS plasmid with the indicated cysteine substitutions in OutD and/or OutB were grown aerobically to allow formation of disulfide bonds in the periplasm. The remaining free thiol groups were then blocked with iodoacetamide and the extent of disulfide bonding was assessed in nonreducing gel, followed by immunoblotting with OutD and OutB antibodies. The same samples were analyzed in reducing gel with 2-mercaptoethanol to estimate the quantities of OutB and OutD. An equivalent amount of cells was loaded into each well. OutB, OutD and their complexes are indicated by an arrow or by an asterisk.

To further map the OutD binding site, in reverse photo cross-linking experiments, *p*BPA substitutions were introduced along the β1 and β3 strands of OutD N0, in place of the residues detected by the MS analysis or spatially close to them (Fig. 6B and C). An obvious OutD-OutB complex reactive with both OutD and OutB antibodies was detected with the OutD_*p*BPA substitutions of E2, S4, S6 (β1), Y42 and M44 (β3) (Fig. 6C). In the OutD N0 structure (Fig. 6B), these residues are located close to each other and hence could interact with the same or proximal sites of OutB HR. Significantly, Y42 and M44 support the OutD cross-linking site identified by the MS analysis (Fig. 6A).

The validity of the identified HR-N0 interaction site was further assessed with *in vivo* disulphide-bonding assay. Several residues of OutB HR that generated a *p*BPA-induced complex were substituted with cysteine and co-expressed with either the wild-type OutD or OutD_Y42C since OutD_M44C variant was barely detectable. Formation of disulphide bonds between spatially proximal cysteine residues were next assessed during bacterial growth. Among the tested combinations, OutB_S139C/OutD_Y42C and OutB_Y137C/OutD_Y42C pairs generated an abundant OutB-OutD complex supporting the proximity of these residues in the functional T2SS (Fig. 6D). Therefore, in spite of the different hydrophobicity of the substituted residues, *p*BPA versus cysteine, and the length of the respective cross-linking, 3 Å versus 7 Å, both *in vivo* approaches confirm the MS data and clearly map one OutB-OutD interacting site to S139 of OutB HR and to a short zone around Y42 and M44 of OutD N0.

### Model of OutD N0-OutB HR interaction

Based on the data of *in vivo* cross-linking experiments, we generated an OutB HR/OutD N0 model by employing the HDOCK server (81). In this docking modelling, the proximity of the residues Y137, S139, R152, R154, Q170 of OutB HR and E2, S4, S6, Y42, M44 of OutD N0 was given as preferable constraints for interacting residues. Among several generated OutB HR/OutD N0 models, one being distinguished by an excellent docking energy of -149.45 and several convincing structural features was retained (Fig. 7A). Firstly, in this model, all the interacting residues of OutB HR and OutD N0, listed above, are placed in a close vicinity. Secondly, a striking aspect of this model is the complementation of several β-strands between the OutB HR and OutD N0 domains. Precisely, the β1-strand of OutB HR defined by the residues S133 to Y137 determines an antiparallel β-sheet with the β1-strand of OutD N0 containing residues S4 to K8; this leads to a complementary five stranded anti-parallel β-sheet implicating the two distinct domains (Fig. 7A). Such a mixed HR/N0 β-sheet is consistent with the even/odd alternation of photo cross-linking patterns observed with the residues of β1-strand of OutB HR and β1-strand of OutD N0 (Fig. 5B and 6C). It seems plausible that formation of this mixed β-sheet together with a rather large HR/N0 interdomain interface could explain high affinity of the OutB HR/OutD N0 interaction observed both *in vivo* and *in vitro* (Fig. 5B, S3 and S12).

**Figure 7.**
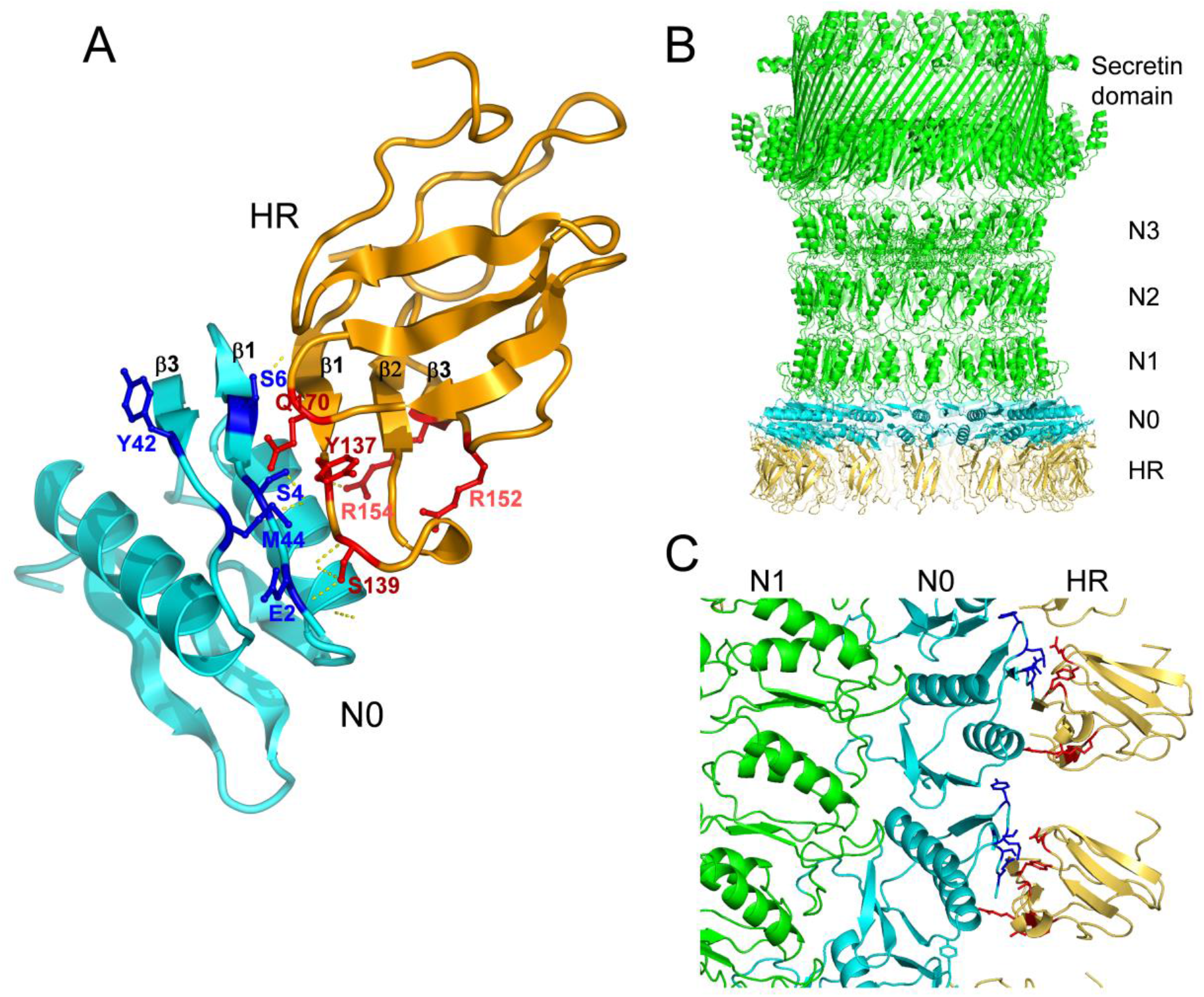
Model of the OutB HR/OutD N0 complex of *D. dadantii*. **A**. OutB HR/OutD N0 interdomain interface modelled according to the *in vivo* cross-linking data. **B**. Model of the 15-meric secretin channel in complex with OutB HR domains, assuming all N0 domains are occupied, front view. **C**. Close up view of the interacting regions between OutB HR and OutD N0; two adjacent N0/HR complexes are shown. OutB HR domains are in gold, OutD is in green with N0 domains in cyan.

The *D. dadantii* OutD is highly similar (66% sequence identity) to the secretin PulD of *K. pneumoniae* whose pentadecameric structure was recently solved by cryo-EM (27). The N0 domains are notoriously flexible, but have been imaged in this high-resolution structure, as a tightly packed ring stabilized in the presence of PulC (PDB 6HCG). Based on PulD structure we generated the full-length *D. dadantii* secretin model, to validate its ability to bind OutB in such a multimeric conformation. The generated model confirms that OutB is indeed able to bind to the 15-meric *D. dadantii* secretin with no clashes (Fig. 7B and C). The steric hindrance of the HR/N0 interface area allows the binding of one copy of OutB to one OutD subunit. Our protein/protein docking model between OutD and OutB HR shows that it can be repeated 15 times to form a complete ring with a relative stoichiometry of OutB-OutD complex of 1:1. In this arrangement, OutB HR domains are located radially inward of the N0 ring without any lateral steric contact clashes with neighboring HR domains, forming a cap to the entry of pentadecameric secretin channel.

### In the functional T2SS, OutC could compete with OutB for the secretin OutD

The HR domain of GspC/OutC components has also been reported to interact with the N0 domain of secretin (31, 32). Interestingly, the OutB HR/OutD N0 complex, that we modelled based on *in vivo* cross-linking data (Fig. 7A), is similar to the crystal structure of GspC HR/GspD N0-N1 complex from *E. coli* (PDB entry 3OSS) (31). Particularly, when the N0 domains in these two structures were superimposed, the corresponding HR domains are also well superposed, such as equivalent secondary structure elements are placed in the two HR/N0 interdomain interfaces (Fig. S11). It seems therefore plausible that in the secretion system, OutB and OutC could also take such equivalent arrangements competing for the secretin. To test this hypothesis, NMR HQSC titration experiments were performed with the individual OutB HR, OutC HR and OutD N0 domains (Fig. S12). In these conditions, OutC HR did not compete with OutB HR for OutD N0 indicating that *in vitro*, OutB HR binds to OutD N0 domain with higher affinity than OutC HR. To assess whether these two HR domains compete for OutD *in vivo*, OutB_R154*p*BPA was expressed, either with or without OutC, from CDBS or DBS plasmid, respectively. In *E. coli*, similar amounts of OutB-OutD complex were generated from these two plasmids (Fig. S13), indicating that OutC does not affect the OutB-OutD interaction in this background. In contrast, in *D. dadantii*, the amount of the generated OutB-OutD complex was reproductively lower in the presence of OutC (Fig. S8 and S13), indicating that within the functional T2SS, OutC could compete with OutB for the secretin binding.

### GspB phylogeny

Search against the UniProtKB database by using the OutB HR domain as template shows the occurrence of GspB homologs in several groups of γ and β-proteobacteria. All these proteins carry a single transmembrane segment and a periplasmic HR domain but vary substantially in length and composition, so that a confident phylogenetic analysis could only be performed with the HR domains of these proteins (Fig. S14). Systematic inspection of *gspB* synteny and domain organization allow us to classify these proteins into seven archetypes (Fig. 2B). In most bacteria, *gspB* gene is co-located with *gspA*. Typically, GspA consists of a cytoplasmic AAA+ ATPase domain fused to a C-terminal transmembrane segment that is followed by a large periplasmic region with a cysteine peptidase-like and a PG-binding domain, such as in *Vibrio* and *Aeromonas* (Fig. 2B). Yet, in *Alteromonas* and *Pseudoalteromonas* groups, GspAs lack the periplasmic region and the cognate GspBs are notably longer than in the other bacteria (Fig. 2B). In addition, GspBs of *Alteromonas* group possess a supplementary cytoplasmic domain of DUF3391 family. Interestingly, in these bacteria, a gene coding for a DUF3391 domain protein is present next to *gspB* (Fig. 2B) indicating a possible origin of DUF3391 domain in GspB. In some bacteria, such as *V. vulnificus*, GspA is fused to the periplasmic portion of GspB in a single GspA-B polypeptide (82). Similar GspA-B fusions of various lengths and domain contents seem to have appeared independently in several groups of γ-proteobacteria (Fig. S14), supporting close functional relationship between GspA and GspB. *E. coli* K-12 and some other close enterobacteria possess an archetypal multidomain GspA in pair with a remarkably small GspB that has no HR domain but carries instead a variable polyampholyte sequence (Fig. 2B). Finally, *Dickeya, Pectobacterium, Klebsiella* and a few other close bacteria lack any *gspA* homolog, and *gspB* gene is co-located with a gene of the OutS/PulS pilotin family. Therefore, depending on bacteria, GspB proteins vary substantially in length and domain composition and act in concert with either a cognate GspA or a pilotin GspS.

### OutB can be efficiently cross-linked to the PG *in vivo*

Separation of *D. dadantii* membrane vesicles in sucrose gradient suggested that OutB is anchored to the PG. Notably, degradation of the PG by lysozyme affects the location of OutB (Fig. 3C and D). In *Aeromonas* and *Vibrio*, GspA carries a specialized PG-binding domain (Fig. 2B) and GspA/GspB complex is involved in the insertion of cognate secretins into or through the PG mesh (57, 83). In *Dickeya* and related bacteria that lack a GspA homolog, OutB has an additional C-terminal polyampholyte sequence (Fig. 2). Therefore, we investigated if in *D. dadantii*, the absence of GspA could be compensated by this C-terminal extension.

The thiol-cleavable cross-linker 3, 3′-dithio-bis (sulfosuccinimidyl) propionate (DTSSP) was used to explore OutB-peptidoglycan interaction. *D. dadantii ΔoutB* cells harboring a plasmid with *outB*, or *outBΔcte*, or empty vector were treated with DTSSP and PG was extracted by SDS-boiling procedure. The PG-bound proteins were next eluted with 2-mercaptoethanol and analyzed by immunoblotting (Fig. 8A). The PG-associated protein OmpA, used as a positive control, was equally eluted from all three PG preparations. In contrast, only OutB but not OutB_*Δ*CTE_ was massively eluted from the PG. Therefore, OutB was efficiently cross-linked by DTSSP to the PG via its C-terminal extension suggesting that OutB CTE is proximal to the PG layer and could interact with it.

**Figure 8.**
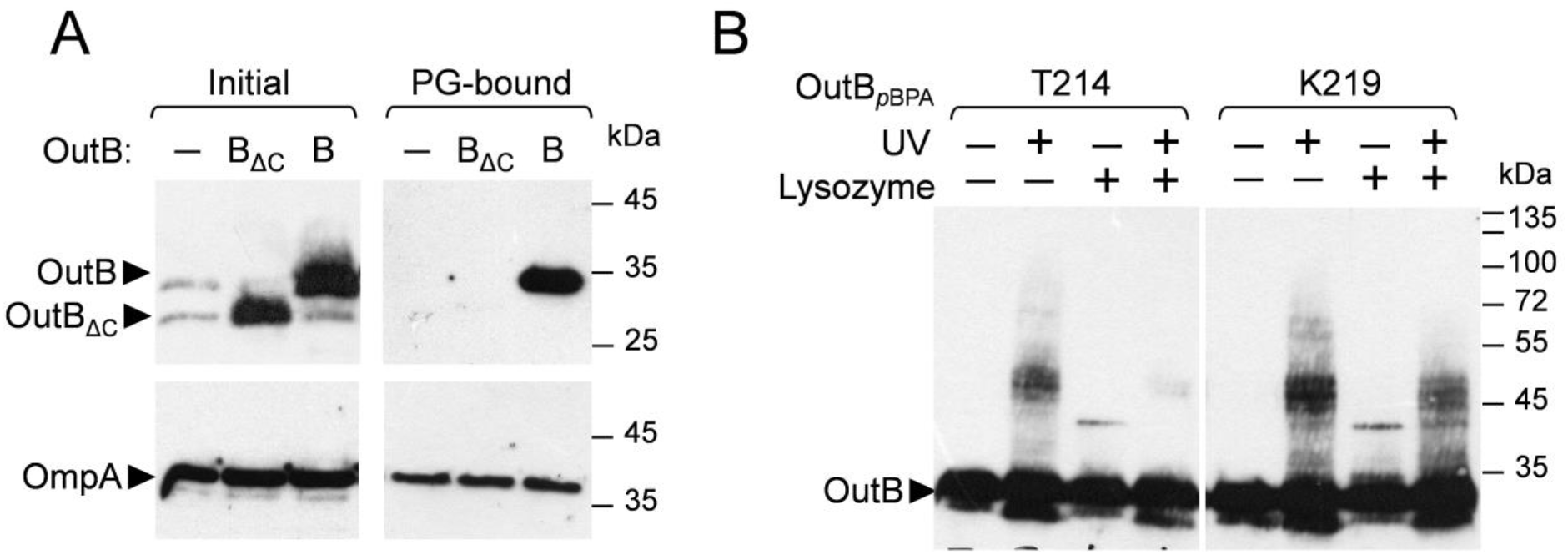
OutB interacts with peptidoglycan. **A**. *In vivo* chemical crosslinking with DTSSP. *D. dadantii outB* A5719 cells carrying either an empty vector, or a plasmid with *outB*, or *outBΔcte* (-, B and B_*Δ*C_, respectively, in ‘Initial’) were treated with DTSSP. The PG was next extracted by SDS-boiling procedure and PG-bound proteins were eluted with 2-mercaptoethanol and analyzed by immunoblotting with anti-OutB or anti-OmpA (’PG-bound’). 25-fold less material, in cell equivalent, was loaded in ‘Initial’ in comparison with ‘PG-bound’. **B**. *In vivo* site specific photo-crosslinking. OutB *p*BPA substitutions (T214 and K219) were expressed in *D. dadantii outB* A5719. Cells were irradiated by UV, treated or not with lysozyme and analyzed by immunoblotting with anti-OutB.

To further explore the OutB-PG interaction, photo cross-linking with *p*BPA, which has a shorter arm length than DTSSP (3 Å *vs* 12 Å) and a shorter time-lapse (3 min *vs* 20 min) was employed. *p*BPA substitutions were introduced into OutB CTE and their cross-linking patterns were assessed in *D. dadantii* (Fig. 8B). Diffuse multi-band adducts of about 40 to 50 kDa over a less-intense ladder-like background was observed. Treatment of cross-linked cells with lysozyme significantly decreased the amount of diffuse adduct and reduced the ladder-like background indicating that the PG-derived sugars are part of these complexes. Taken together these data suggest that *in vivo*, the C-terminal extension of OutB interacts with the PG. The functional relevance of the CTE was determined using the *D. dadantii outBΔcte* mutant, where the reduced pectinase and cellulase halos indicate diminished secretion (Fig. 1B). The effect was not as substantial as the complete *outB* deletion, but was significant, revealing the functional importance of the C-terminal extension of OutB.

## Discussion

Here we show that OutB is essential for effective assembly and function of the *D. dadantii* T2SS and for expression of full bacterial virulence. We reveal that OutB has three functions linked to distinct regions of the protein: 1) The N-terminal transmembrane helix anchors OutB to the IM; 2) The conserved periplasmic domain of OutB (HR) binds tightly the N0 domain of the secretin OutD and assists its targeting to the outer-membrane, and 3) The C-terminal region (CTE) attaches OutB to the peptidoglycan, anchoring the entire secretion system to the bacterial cell wall.

The crystal structure reveals that the conserved periplasmic domain of OutB has the same fold as the homology regions (HR) of the T2SS GspC and the T4P PilP components (31, 74-76). In spite of low sequence conservation (10 to 17% of identical residues), OutB HR structure is highly similar to these HR domains (r.m.s.d. 1.7 Å to 2.8 Å for 58 equivalent CA atoms). This structural similarity coincides with the fact that all these HR domains interact with the N0 domain of cognate secretins (31, 32, 74, 84). By combining structural, *in vivo* and modelling approaches, we unravelled the molecular details of the OutB HR/OutD N0 interaction. Particularly, site-specific photo cross-linking followed by mass spectrometry analysis allowed an unambiguous identification of the OutB-OutD interaction site *in vivo*. The validity of this site was further confirmed with reverse photo cross-linking and *in vivo* disulphide bonding. Consequently, the OutB HR/OutD N0 model constructed after these data reflects a functionally relevant protein contact within the functional secretion system. NMR spectroscopy is roughly consistent with the cross-linking experiments but for various technical reasons failed to give a comprehensive list of residues involved in the OutB HR/OutD N0 interface.

The N0 domains constitute the entry of the T2SS secretin channel, which controls the recruitment of substrates and communicates with the IM components of the secretion machinery (29-32). Recent cryo-EM tomography of the *L. pneumophila* T2SS showed high flexibility of the N-terminal portion of the secretin channel that could reflect different functional states and interactions with various partners (34). The occurrence of two components, OutB and OutC, carrying structurally similar HR domains that interact with the same N0 domain of the cognate secretin, raises the questions of the hierarchy of these interactions within the T2SS. Noteworthy, the fact that OutB HR/OutD N0 interface modelled after our photo-CL data is strikingly similar to that of the crystal GspC HR/GspD N0-N1 complex (Fig. S11) (31) suggests that these two interactions are spatially incompatible and hence could not occur simultaneously. NMR experiments show that *in vitro* OutB HR binds OutD N0 more tightly than OutC HR does, with the affinities ranging in 1 mM and 10 µM order for OutC and OutB, respectively. (Fig. S12). Similarly, in *E. coli* carrying a partially reconstituted transmembrane complex, OutC does not affect the OutB-OutD complex (Fig. S13). However, in *D. dadantii*, OutC seems to interfere with the OutB-OutD interaction (Fig. S8 and S13). Since GspC/OutC components are involved in substrate recruitment (29, 30), it is tempting to speculate that OutC-OutD interactions are prevalent during the course of secretion while OutB-OutD interactions play a role in the assembly, maintenance and scaffolding of the secretin channel.

We present a model of the OutD N0/OutB HR interaction in the context of the 15-meric secretin channel (Fig. 7B). In this model, each OutB HR binds one OutD N0 domain in 1:1 ratio, such as OutB HR domains are located radially inward of the N0 ring with no lateral contacts with neighboring HR domains. The model presented here is therefore rather symmetric and constrained compared to the more open and dynamic arrangement, as it can be anticipated in the functional secretion system. Indeed, the functional oligomeric state of the secretin channel within the functional T2SS has not been ultimately determined. Recent cryo-EM studies revealed a C15-fold symmetry of the T2SS secretins (15, 21, 23, 28), while *in vivo* disulfide-bonding analyses are consistent with a C6 symmetry of a hexamer of dimers (32, 71). In agreement with the latter, the N-terminal portion of *P. aeruginosa* secretin XcpQ, comprising the N0-N1-N2 domains, self-assemblies into a hexamer of dimers, forming *in vitro* C6 symmetry ring-shaped channels (85). Lithgow and colleagues proposed an elegant solution to this apparent discrepancy (23) suggesting that the C15 symmetry observed in the secretin and N3 domains is further followed by the pseudo 6-fold symmetry for the N0-N1-N2 domains, compatible with C12 or C6 symmetry of the assembly platform, either hexameric as GspE, GspL and GspM, or dodecameric, as GspC (27). The common symmetry element between C6, C12 and C15 symmetries is a three-fold axis. Therefore, C12 and C15 complexes could have common three-fold symmetry if every fifth N0 domain is unoccupied. Yet it is not clear how such an arrangement would be enforced; thus it is plausible that the complex overall is asymmetric unless three-fold symmetry is imposed in the periplasm by other components.

The role that OutB plays to maintain the secretin channels is closely linked to its ability to bind peptidoglycan. Chemical cross-linking experiments with DTSSP and *p*BPA showed that the C-terminal extension of OutB interacts with the peptidoglycan. The PG is composed of glycan chains connected by short peptides that together form a mesh surrounding bacterial cell. In proteobacteria, the PG layer is located in the periplasm and it is attached to the OM through a covalent linkage to the Braun’s lipoprotein Lpp (86). Several multi-protein structures and machineries span the entire cell envelope. The size of the peptidoglycan mesh (about 2 nm) is not compatible with the external dimensions of these large complexes and specialized or housekeeping transglycosylases are recruited for the local rearrangement of the peptidoglycan layer in the course of their assembly through the cell envelope (87, 88). The attachment of these transenvelope machineries to the PG mesh is usually ensured by the specialized PG-binding modules. For instance, the T4P system possesses two components carrying specialized PG-binding domains, secretin PilQ and TsaP, showing the importance of a solid attachment to the PG (25, 52, 53). Generally, the PG-binding components represent a less conserved part of respective transenvelope systems suggesting that anchorage to the PG has evolved and diversified more recently, in accordance with the particular cell wall context (55). For instance, *in silico* analysis of the T6SSs has revealed surprising richness and variability of the associated PG-binding modules (89). The authors have proposed the notion of more or less “evolved” PG-binding modules that have been developed from a few ancestral proteins by gene fusion and loss of some nonessential regions.

Phylogenetic analysis shows that in many bacteria, such as in *Vibrio* and *Aeromonas*, GspB acts in concert with a multi-domain ATPase GspA that carries a specialized PG-binding domain, (Fig. 2B and S14). In this group, some “more evolved” GspA and GspB are naturally fused into a single polypeptide (59). A C-terminal extension, CTE, similar to that of OutB is only present in the *Dickeya*/*Klebsiella* group that lacks a GspA counterpart (Fig. 2). We showed that the OutB CTE could interact with peptidoglycan. Therefore, it is plausible that in these GspBs, the absence of the PG-binding partner, GspA, is compensated by a polyampholyte C-terminal sequence that interacts with PG. The molecular mechanism of this interaction remains unclear since the OutB CTE does not correspond to any known PG-binding domain. However, the residue combinations of OutB CTE are reminiscent of those in some sugar binding lectin domains. For instance, the bactericidal C-type lectins of RegIII family recognize the PG carbohydrate moiety via an EPN-like motif located at an extended loop region and the variations of this motif, EPN, QPD, EPS, etc. alter its sugar binding specificity (90). It is possible that the OutB C-terminal extension could bind peptidoglycan in a similar way. Some other bacterial proteins, apparently lacking a specialized PG-binding domain, have been shown to bind the PG, e.g. the pilotin InvH from the T3SS of *Salmonella enteritica* (91). Interestingly, in *L. pneumophila* T2SS that lacks any GspB or GspA homolog, the secretin LspD itself carries a SPOR-like PG-binding domain at the N-terminus, prior to the N0 domain (34). Thus, the modes of attachment of T2SS secretins to the PG vary in different bacteria.

In *Dickeya*/*Klebsiella* group, *gspB* is co-located with the gene coding for GspS pilotin (Fig. 2B). We propose a model wherein OutB and OutS play complementary functions in targeting, assembly and scaffolding of the secretin (Fig. 9). OutB could act early during OutD export by binding the N0 domain, the first part of the secretin released into the periplasm. In this way, OutB could orient the secretin and prevent its misinsertion into the IM. In the later step, pilotin binding to the C-terminal S-domain, with the assistance of the Lol machinery, would target and anchor the C-terminal portion of the secretin into the OM (36), while the N-terminal entry of secretin pore remains connected to the IM via OutB. At the same time, the C-terminal extension of OutB would attach the secretin to the PG. Therefore, GspB/OutB has a major role in the assembly of the secretin channel providing it scaffolding with the inner membrane assembly platform and the peptidoglycan layer.

**Figure 9.**
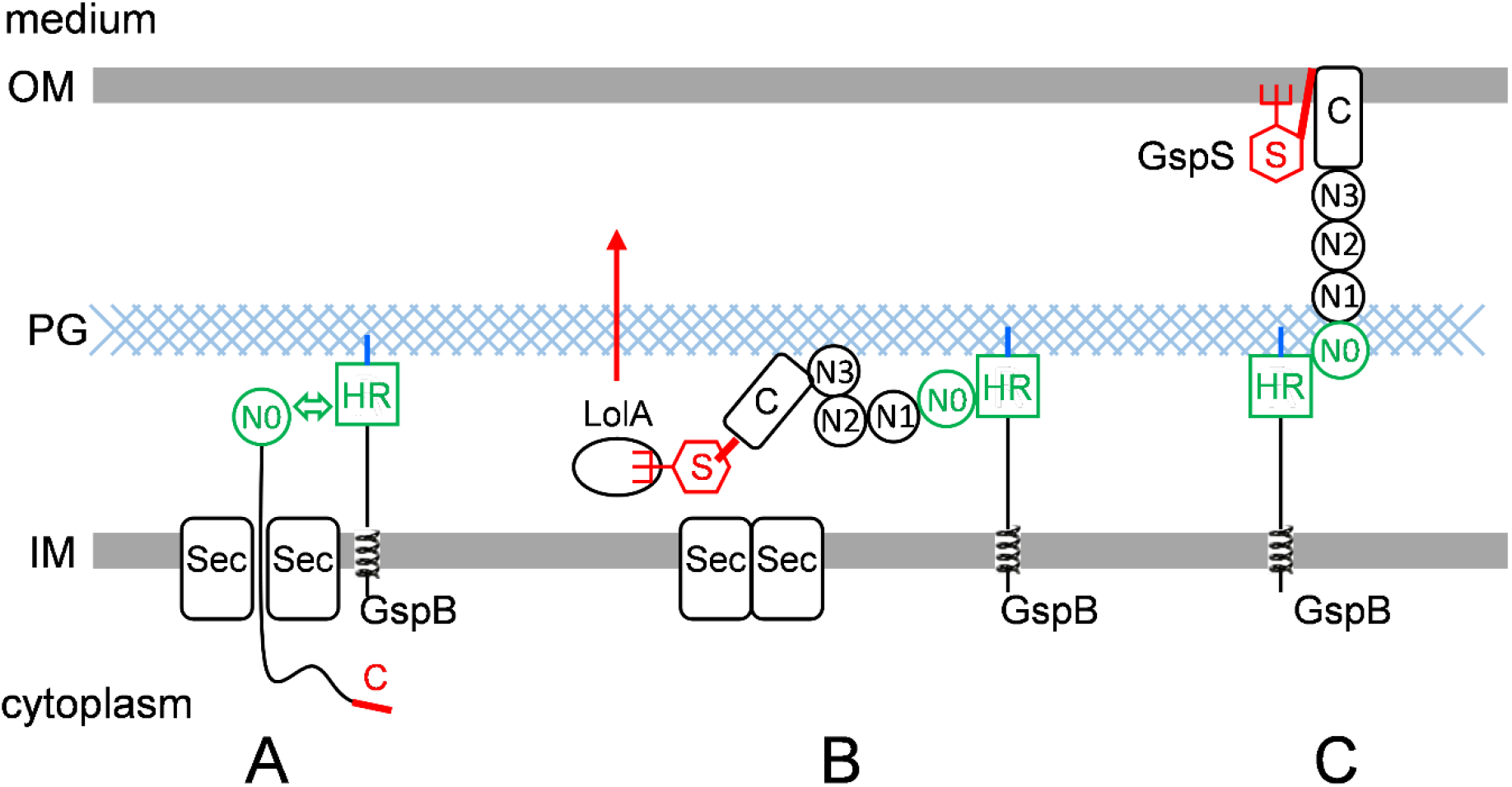
OutB and OutS play complementary functions in targeting, assembly and scaffolding of secretin. **A**. OutB acts early during OutD export by binding the N0 domain, the first part of the secretin releasing into the periplasm. In this way, OutB could orient the secretin and prevent its mis-insertion into the IM. **B**. In the later step, pilotin OutS (red hexagon) binds the C-terminal S-domain (red segment) and, with the assistance of the Lol machinery, targets the C-terminal part of the secretin into the OM where several secretin subunits form a channel. **C**. OutB binds to and stabilizes the N-terminal entry of the secretin channel providing scaffolding with the IM and the PG layer. For simplicity, only one subunit of each protein is shown.

## Materials and Methods

### Strains, plasmids and growth conditions

The bacterial strains and plasmids used in this study are listed in Tables S3 and S4. *D. dadantii* mutant strains carrying chromosomal *outB* mutant alleles were constructed by marker exchange-eviction mutagenesis or by homologous recombination as described (92). The bacteria were grown in Luria-Bertani (LB) or in M9 minimal medium at 28°C with shaking at 120 rpm. If necessary, glucose, glycerol or galacturonate were added at 0.2 % and antibiotics were added at the following final concentrations: ampicillin, 100 µg/ml^-1^; kanamycin, 50 µg/ml^-1^ and chloramphenicol, 25 µg/ml^-1^. DNA manipulations were carried out using standard methods. Site-directed mutagenesis was performed with the PrimeSTAR Max DNA Polymerase (TaKaRa) and the primers are listed in Table S5. The sequences of mutant and amplified genes were checked (Eurofins MWG Operon).

### Protein purification

DNA sequence encoding the conserved periplasmic domain of *D. dadantii* OutB was cloned into the pGEX-6P-3 vector (GE Healthcare) encoding a cleavable N-terminal Glutathione S-transferase (GST) affinity tag (Table S4). Three GST-OutB constructs encoding OutB residues 112-220, 112-202 and 112-192 were expressed in *E. coli* BL21 (DE3) cells (Stratagene). The bacteria were grown in Luria broth supplemented with 100 µg/ml ampicillin to an OD_600_ of 0.6 next, induced with 1 mM isopropyl-β-D-thiogalactopyranoside (IPTG) and grown for an additional 16 h at 20°C. The cells were suspended in phosphate buffered saline (PBS) containing 0.01 M phosphate buffer, 0.0027 M KCl and 0.137 M NaCl, pH 7.4 and lysed by sonication. Cell debris was removed by centrifugation at 35,000g for 20 min and the supernatant from 1 l of cell culture was incubated with 2.5 ml of Glutathione Sepharose beads (GE Healthcare) at 4°C for 6 h The Glutathione Sepharose beads were then collected in a PD10 column and washed with 50 ml of PBS. GST tag was on column cleaved by addition of 20 µl of PreScission protease to 5 ml of Glutathione Sepharose in buffer containing 50 mM Tris-HCl, 150 mM NaCl, 1 mM EDTA, 1 mM DTT, pH 7.0 at 4°C for 6 h. N0 and N0-N1-N2 domains of OutD (residues 1 to 85 and 1 to 258 of signal peptide-less OutD, respectively, Table S4) were produced in *E. coli* BL21(DE3) and purified by nickel-affinity chromatography as described previously (33). For NMR spectroscopy, uniformly ^15^ N- and ^13^ C-labeled proteins were produced by growing cell cultures in M9 minimal medium that contained 1 g/l ^15^ N-ammonium chloride and 2 g/l ^13^ C-D-glucose (Cambridge Isotope Laboratories, Inc.) as the sources of nitrogen and carbon, respectively. Prior to crystallization trials and NMR spectroscopy, the purified proteins were run on Superdex S200 size-exclusion column and concentrated with Vivaspin devices (Sartorius).

### Crystallization

Crystallization conditions were screened using hanging drop vapour diffusion in 96 well plates set up with a nanodrop Mosquito liquid handling robot (TTP Labtech). Protein and precipitant drops were in the range of 200-400 nl. OutB (residues 112 to 202) crystals suitable for X-ray analysis grew in a solution containing 0.2 M MgCl_2_, 0.1 M Tris, 30% w/v PEG4000, pH 8.5 (Molecular Dimensions Structure Screen 1 condition C9). Selected crystals were transferred to a solution containing mother liquor plus 5% propane-1,2-diol as a cryoprotectant and flash cooled in liquid nitrogen. Diffraction data were collected at 100K using a Pilatus 300M detector at beam line I04 at Diamond Light Source (DLS), Oxfordshire. Data were integrated and scaled by XDS through the Xia2 software package (93) available through DLS. The structure was solved by molecular replacement using Phaser from the CCP4 software package (94) using as a search model *E. coli* GspC-HR structure (PDB code 3OSS) detected using PHYRE2 (95). The model was refined using REFMAC v5.8 (96) and 10% of the data were excluded to be used for validation purposes. OutB structure coordinates have been deposited at the Protein Data Bank in Europe (PDBe) database with accession codes 4WFW.

### NMR

NMR spectra were acquired using the shortest OutB HR domain (residues 112-193) at 15°C using Bruker Avance 600 and 700 MHz spectrometers equipped for ^1^ H, ^15^ N, ^13^ C triple resonance experiments. All spectra were processed using NMRPipe/NMRDraw (97) and analyzed using XEASY (98). HNCA, HN(CO)CA, HNCO, HNCACB, and CBCA(CO)NH experiments were used to obtain sequence-specific ^1^ HN, ^15^ N, ^13^ Cα, ^13^ Cβ, and ^13^ C’ backbone assignments. The NMR samples had a typical concentration of 0.2 mM in 20 mm Tris-HCl, pH 7.0, and 10% D_2_O. NMR titration was achieved by recording a ^1^ H-^15^ N HSQC/ ^1^ H-^15^ N Sofast-HMQC spectra of the labeled protein. Cross titrations were used to eliminate the dilution of the labeled protein. In cross titration, two samples were prepared, one sample was labeled protein only and the second sample had the same concentration of labeled protein and a ten equivalents of unlabeled titration partner. 150 mM NaCl was included in the cross-titration studies.

### Structure modeling

A homology modelling of the *D. dadantii* OutD N0 domain (residues 1 to 78 of the matured, signal peptide-less OutD, UniProtKB entry Q01565) was conducted using the trRosetta server (99) with restraints from both deep learning and homologous templates. The confidence of the predicted model is optimal with a TM-score of 0.912 on a maximum of 1.0. Using this OutD N0 model, we then performed a protein/protein docking with the HR domain of OutB (PDB entry 4WFW). In this order, the HDOCK server (81) was employed with the contacting residues, identified by the photo cross-linking experiments, given as preferable constraints (namely Y137, S139, R152, R154, Q170 of OutB HR and E2, S4, S6, Y42, M44 of OutD N0). In addition, the docking algorithm identified PDB entry 3OSS as template (the crystal structure of enterotoxigenic *E. coli* GspC-GspD complex). This gave results to several distinct models, one being distinguished by an excellent docking energy of -149.45 and several convincing structural features. This last was retained for subsequent steps.

A homology modelling of the full-length OutD (residues 1 to 683) was then produced using again the trRosetta server (99). A good model with a TM-score of 0.597 was produced. We then reconstructed the full-length OutD/OutB HR domain interface using the previously identified protein/protein docking pose. Finally, this OutD/OutB HR model was employed to reconstruct the *D. dadantii* secretin pentadecameric model. In this goal, we used the cryo-EM structure of the *K. pneumoniae* type II secretion system outer membrane complex (PDB entry 6HCG) as template (27). We superimposed the OutD/OutB HR oligomer model to each of the 15 units of PulD with PyMOL v1.3 (The PyMOL Molecular Graphics System, Schrödinger, LLC) using its ‘cealign’ function which is very robust for 3D structural alignments of proteins with little to no sequence similarity. We therefore obtained the 15mer model of OutD/OutB HR and we concluded that no further minimization step was needed as its visual inspection demonstrated that no clashes were present in the N-terminal area harboring the OutD/OutB HR interface.

### Pull-down assays

OutB domains of interest fused to GST were co-expressed with various combinations of N0 to N3 domains of OutD from the same plasmid (Table S4) in *E. coli* BL21(DE3) and co-purified with batch method as described (100). Briefly, the cells were broken in a French press in TSE buffer (50 mM Tris-HCl, 100 mM NaCl, 1 mM EDTA, pH 7.5), supplemented with 0.1% (v/v) Triton X-100 and incubated with Glutathione Agarose (Machery Nagel). Non-bound proteins were washed away for six times with the same buffer and the bound proteins were eluted with Laemmli sample buffer, separated by SDS-PAGE and stained with Coomassie or probed by immunoblotting with anti-OutD antibodies.

### Site-specific in vivo photo cross-linking

*D. dadantii* or *E. coli* MG1655 cells, carrying pSup-BpaRS-6TRN (77), pREP4 (Qiagen) and a DBS-derived plasmid carrying an *outB_TAG* or *outD_TAG* substitution were grown in BL supplemented with ampicillin, kanamycin and chloramphenicol at 28°C for 16 h. 1 to 1.2 ml cultures were then collected, washed in M9 and diluted (∼three-fold) to an OD_600_ of 0.5 into M9 medium containing 1 mM *p*BPA, 0.2% glycerol, 0.01% casamino acids and appropriate antibiotics. To induce synthesis of pectinases and Out proteins, *D. dadantii* cultures were additionally supplemented with 0.2% galacturonate. After a 1 h of growth, isopropyl-β-D-thiogalactopyranoside (IPTG) was added to 1 mM and the cultures were followed for an additional 3 h or 5 h for *E. coli* and *D. dadantii*, respectively. Afterward, a 1.4 ml portion of cells was either chilled on ice (control) or placed in a glass Petri dish and irradiated with an UV lamp (Bio-Link BLX model, 365 nm) for 3 min at 10 cm distance. The cells were chilled on ice, collected by centrifugation, lysed in Laemmli sample buffer and resolved on 9% SDS PAGE. The proteins were detected by immunoblotting using antibodies raised against OutB or OutD.

### Purification of OutB-OutD complex for mass spectrometry analysis

*E. coli* MG1655 cells, carrying pSup-BpaRS-6TRN (77), pREP4 (Qiagen) and DBS-derived plasmid co-expressing an *outB_TAG* variant together with *outD_Strep* were grown and subjected to photo cross-linking exactly as described in the chapter above. Precisely, a series of 10 tubes with 5 ml culture was grown and exposed together to UV in nine 10 cm diameter Petri dishes in the Bio-Link BLX chamber. After 3 min irradiation with UV at 365 nm, the cells were chilled on ice, collected by centrifugation and stored at -20°C. The frozen cells from each series (50 ml culture) were resuspended in 4 ml TSE buffer (50 mM Tris-HCl pH 8.0, 100 mM NaCl, 1 mM EDTA, 1 mM PMSF) supplemented with 1 mg/ml lysozyme and incubated for 15 min. The cell suspension was next supplemented with SDS to 1% and boiled for an additional 15 min. Non-soluble cell material was removed by centrifugation at 7,000g for 10 min and supernatant was ten-fold diluted with TSE buffer supplemented with 1% Triton X-100 and incubated with 2 ml of Step-Tactin XT Superflow (IBA) resin for 2 h at 15°C. The resin was next washed five times with 10 ml of the same buffer and OutD-strep protein and complexes were eluted with TSE buffer supplemented with 1% Triton X-100 and 50 mM biotin. The eluted proteins were precipitated with five volumes of ethanol, resuspended in Laemmli sample buffer and separated on 9% SDS-PAGE.

### Liquid chromatography tandem mass spectrometry of OutB_pBPA-OutD complexes

The protein bands corresponding to OutB_*p*BPA-OutD complexes were excised from a stained gel and subjected to digestion with chymotrypsin and trypsin. Precisely, the gel pieces were destained with a 50% (v/v) mixture of 50 mM ammonium bicarbonate in acetonitrile, reduced with 10 mM DTT (56°C, 1 h) and alkylated with 55 mM iodacetamide (25°C, 45 min, at darkness). Proteins were first digested with chymotrypsin (Promega) (12.5 ng/µl, 25°C overnight). The 1/10 of the sample was kept for injection and the 9/10 were digested with trypsin (Promega) (12.5 ng/µl, 37°C overnight). After quenching with formic acid (FA) to 5%, the two digests were analyzed with an Ultimate 3000 nano-RSLC (Thermo Fischer Scientific) coupled on line with a Q Exactive HF mass spectrometer via a nano-electrospray ionization source (Thermo Fischer Scientific).

The samples were loaded on a Acclaim PepMap100 C18 trap column (20 × 0.075 mm, 100 Å pore size) (Thermo Fischer Scientific) for 3 min at 5 µl/min with 2% acetonitrile, 0.05% trifluoracetic acid (TFA) in H_2_O and then separated on a Acclaim PepMap 100 C18 analytical column (50 × 0.075 mm, 100 Å pore size) (Thermo Fischer Scientific) linear gradient of A (H_2_O, 0.1% FA) and B (100% acetonitrile, 0.1% FA) solvents. Settings were: initial conditions of 4% to 50% of B in 60 min then from 50% to 95% of B in 2 min, hold for 10 min and returned to the initial conditions in 1 min for 14 min. The total run duration was set to 90 min at a flow rate of 300 nl/min. The oven temperature was kept at 40°C.

### Mass spectrometry data processing

Samples were analyzed with a TOP20 HCD method. MS data were acquired in a data dependent strategy selecting the fragmentation events based on the 20 most abundant precursor ions in the survey scan (300-1600 Th). The resolution of the survey scan was 60,000 at m/z 200 Th and for MS/MS scan the resolution was set to 15,000 at m/z 200 Th. The Ion Target Value for the survey scans in the Orbitrap and the MS/MS scan were set to 3E6 and 1E5 respectively and the maximum injection time was set to 60 ms for MS scan and for MS/MS scan. Parameters for acquiring HCD MS/MS spectra were as follows: collision energy of 27 and an isolation width of 2.0 m/z. The precursors with unknown charge state, charge state of 1 and 8 or greater than 8 were excluded. Peptides selected for MS/MS acquisition were then placed on an exclusion list for 20 s using the dynamic exclusion mode to limit duplicate.

Raw data were first submitted to Proteome Discoverer 2.2 using SEQUEST HT search engine. A FASTA file composed of the OutB and OutD sequences was created. Precursor mass tolerance was set at 10 ppm and fragment mass tolerance was set at 0.02 Da, and up to 2 missed cleavages were allowed. Enzyme parameter was set to a combination of trypsin and chymotrypsin. Oxidation (Met), acetylation (Protein N-terminus) and the S^139^ *p*BPA modification delta mass (+164.0626 Da) were set as variable modifications. Carbamidomethylation (Cys) was set as fixed modification. Validation of the identified peptides were done by a ‘Fixed value’ approach based on SEQUEST scores. *p*BPA crosslinked peptides were mapped with the StavroX part of MeroX (2.0.1.4 version http://stavrox.com). Following parameters are set: raw date converted in mgf format; site 1: Ser^139^ BPA (noted as x); site 2: all amino acids with methionine oxidation as variable modification, precursor ion precision at 5 ppm and 0.02 Da on fragments. Validation of the candidates was based on a target-decoy analysis. A False Discovery Rate (FDR) was calculated for each candidate and a FDR score cut-off is applied at the end.

### Chemical cross-linking and PG isolation

Peptidoglycan was extracted from *D. dadantii* cells exponentially grown at BL broth supplemented with ampicillin and galacturonate. The cells (∼5×10^10^) were washed with 50 mM sodium phosphate buffer pH 7.2, resuspended in 2 ml of the same buffer and dropped in 2 ml of boiling SDS 10% solution. After 1 h of boiling, the suspension was stored overnight at 30°C and peptidoglycan was collected by centrifugation at 48,000 rpm in SW 55 Ti rotor (Beckman Coulter) for 2 h at 30°C. Afterward, peptidoglycan was resuspended in 4 ml of 2% SDS solution, boiled again for 30 min and collected by centrifugation at 48,000 rpm in SW 55 Ti rotor for 1 h. Aliquots of purified peptidoglycan were boiled with Laemmli loading buffer containing 2-mercaptoethanol and the eluted PG-bound proteins were analyzed by immunoblotting with anti-OutB and anti-OmpA antibodies.

### Enzymatic assays

Plate assay for pectinase and cellulase secretion was performed as described (92). Bacteria were patched onto polygalacturonate or carboxymethyl cellulose containing plates, for pectinase and cellulase activities respectively, and grown at 28°C for 12 to 20 h. The plates were next flooded with copper acetate or Congo red solutions, respectively, to reveal degradation halos. The halo size reflects the efficiency of pectinase and cellulase secretion. For β-galactosidase activity assays, *E. coli* MC3 carrying pREP4 and a DBS-derived plasmid were grown in LB supplemented with 1 mM IPTG, ampicillin and kanamycin at 28°C for 14 h. Afterward, the cells were permeabilized with toluene and 0.05 % SDS and used in β-galactosidase assay, as described (101).

### Cell fractionation

The inner and outer membrane vesicles were separated on discontinuous sucrose density gradient. *E. coli* MC3/pREP4 cells carrying a DBS derived plasmid were grown in LB medium. 100 ml of LB supplemented with ampicillin and kanamycin was inoculated with non-induced overnight culture to an OD_600_ 0.1 and cultivated at 120 rpm and 28°C for 2 h, then 1 mM IPTG was added and culture followed for 7 h. Cells were collected, resuspended in 5 ml of 50 mM HEPES-NaOH pH 7.4 containing protease inhibitor cocktail (Halt EDTA-free, Thermo) and disrupted by French pressure. Unbroken cells and large debris were eliminated at 5,000g for 10 min. The cell-free extracts were supplemented with sucrose to 30% and loaded on a discontinuous sucrose 35 to 65% gradient (with a 5% step). The gradients were centrifuged at 48,000 rpm in SW 55 Ti rotor (Beckman Coulter) for 70 h, then 0.3 ml fractions were collected from the bottom and analyzed by immunoblotting with the indicated antibodies. *D. dadantii* were grown on polygalacturonate containing plates at 28°C for 36 h (3 plates per strain). The plate-grown cells were collected, washed with HEPES-NaOH pH 7.4, broken by French press and separated as above. When indicated, French pressure cell-free extracts were treated with lysozyme (0.1 mg/ml) at 25°C for 30 min prior loading on sucrose gradient.

### Immunoblotting

The protein extracts were separated by electrophoresis on either 7.5, 9, 12 or 4-20 % SDS-PAG (home-made or Bio-Rad), transferred on Immobilon-P membrane (Millipore) and analyzed by immunoblotting with the polyclonal rabbit antibodies raised against OutB, OutC and OutD (39, 60), TolA (provided by J.C. Lazzaroni, MAP Lab), OmpA (generated against recombinant *D. dadantii* OmpA, this study), PspA (provided by Hendrick Osadnik, Leibniz Universität Hannover) or monoclonal mouse Ro-LPS antibody (provided by J.C. Lazzaroni, MAP Lab).

### Phylogenetic analysis

Candidate GspB homologs were searched at a protein level by using BLAST (102) and OutB HR (residues L127 to P191) as a query sequence against the Bacteria UniProtKB database. 1000 best hits were manually inspected to remove incomplete or fragment sequences thus keeping 936 sequences. Sequences were then clustered with CD-HIT software (103) with 30% identity threshold to keep 118 representative sequence clusters. Multiple sequence alignments were done with T-Coffee program (104) and the HR domain sequences (correspond to residues L127 to P191 of OutB) were selected and used for phylogenetic analyses. Phylogenetic analyses were performed, with the IQ tree web server in auto mode (105). Consensus tree was constructed from 1000 bootstrap trees. Robinson-Foulds distance between ML tree and consensus tree: 14. Phylogenetic tree was visualized with iTOL software (106).

## Acknowledgements

We thank G. Condemine and other team members for critical reading of manuscript. SZ was supported by the China Scholarship Center program. VES was supported by the funding from the Centre National de la Recherche Scientifique (CNRS), ANR program SYNERGY_T2SS ANR-19-CE11-0020 and CNRS PICS 161532 program. SG and RWP are supported by the Biological and Biotechnological Sciences Research Council (BB/M002969/1) and RWP by the Higher Education Funding Council for England and Queen Mary University of London. We thank Dr. Geoff Kelly and Tom Frankiel (Medical Research Council [MRC] Biomedical NMR Centre) for assistance with NMR spectroscopy. NMR data were recorded in the MRC Biomedical NMR Centre at the Francis Crick Institute, which receives core funding from Cancer Research UK Grant FC001029; Medical Research Council Grant FC001029; and Wellcome Trust Grant FC001029. The authors would also like to thank Diamond Light Source for beamtime, and the staff of beamline I04 for assistance with crystal testing and data collection.

