## Supplemental data for "Scaffolding protein GspB/OutB facilitates assembly of the *Dickeya dadantii* type 2 secretion system by anchoring the outer membrane secretin pore to the inner membrane and to the peptidoglycan cell wall"

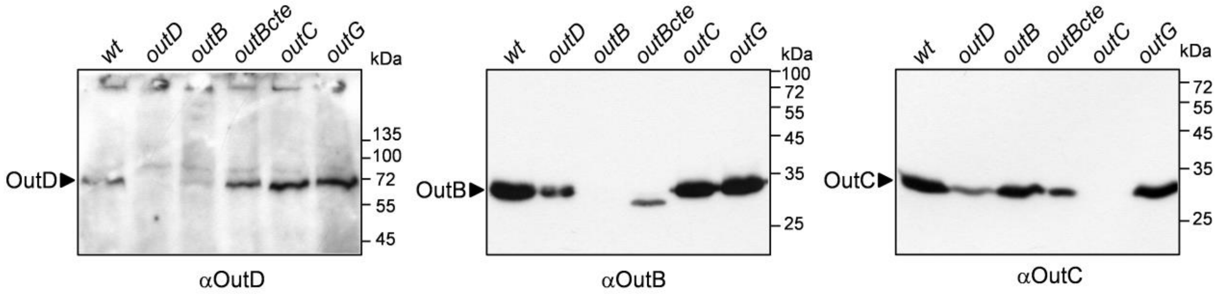


**Figure S1. In *D. dadantii*, *outB* mutation diminishes the amount of OutD.** *D. dadantii* wild type strain, *outD*, *outB, outB*$_{\Delta cte}$, *outC* and *outG* mutants were grown for 14 h at 30°C on the pectin containing medium. An equivalent amount of cells was loaded into each well and probed by immunoblotting with the indicated antibodies. Positions of the Out proteins and molecular mass standards are indicated.

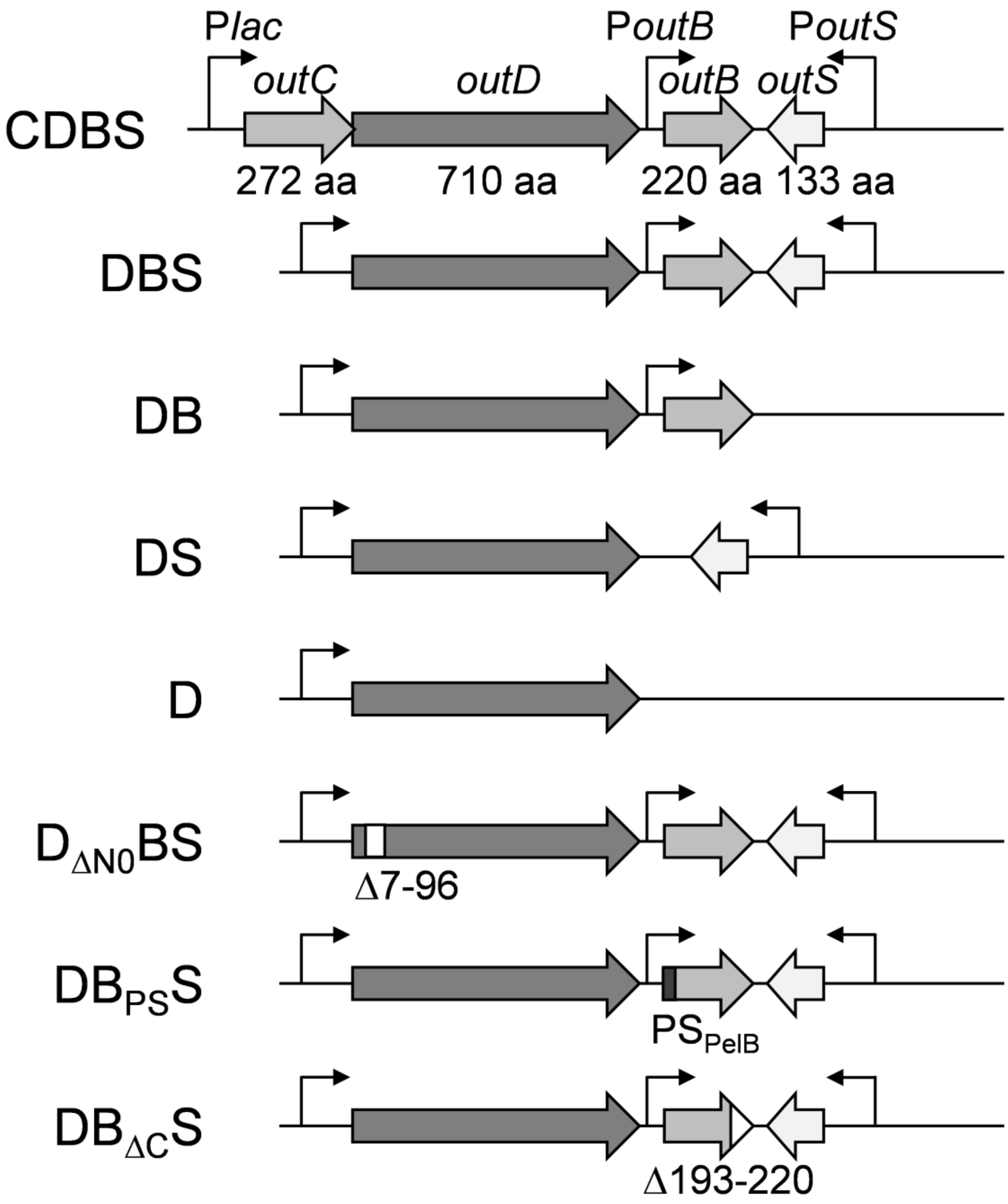


**Figure S2. Schematic of the plasmids used in this study.** The indicated gene combinations were cloned into pGEM-T plasmid (Promega) such as *outC* and *outD* are expressed under the control of P*lac*.

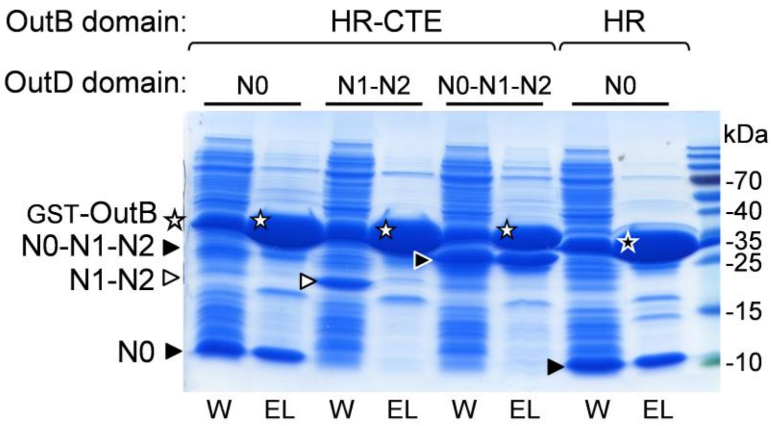


Figure S3. **GST pull-down maps the OutB-OutD interaction to OutB HR and OutD N0 domains.** The C-terminal portion of OutB (HR-CTE or HR alone) was fused to GST and co-expressed in *E. coli* BL21(DE3) together with the N-terminal domains of OutD (indicated at the top). The whole cell extracts (W) were next purified on Glutathione agarose (EL) and analysed by SDS-PAGE. The protein domains and molecular mass standards are indicated.

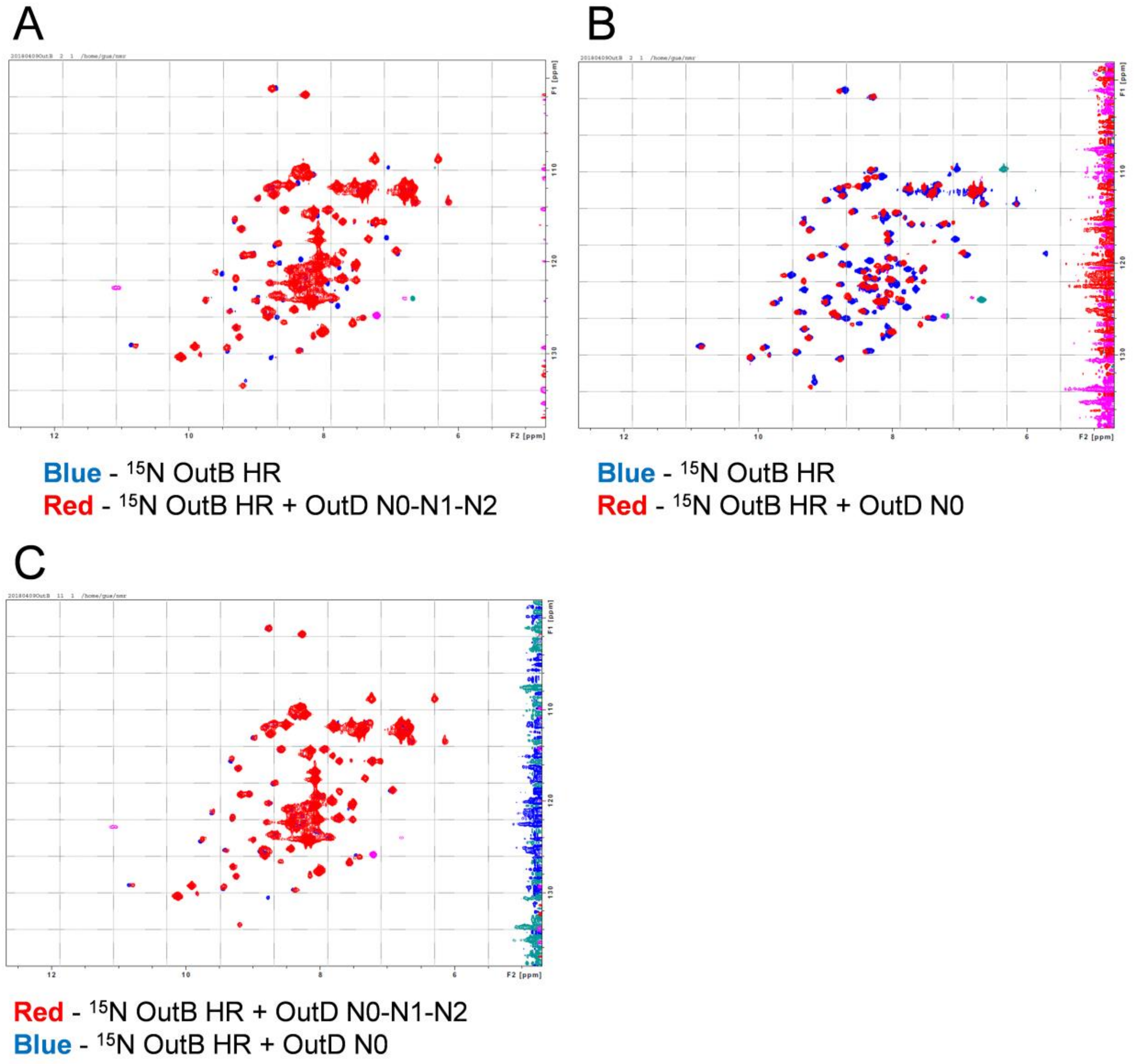


**Figure S4. NMR HQSC experiments show that OutD N0 domain interacts with OutB HR domain. A** and **B**. $^{1}$H- $^{15}$N-HSQC spectra measured with $^{15}$N-labelled OutB HR domain (residues P112-K220), either alone (blue spectra), or (**A**) with unlabeled OutD N0-N1-N2 (residues A1-V258), or (**B**) with unlabeled OutD N0 (residues A1-S85) (red spectra). The concentration of each protein is 200 µM. **C**. The spectra from panel ***A*** (HR + N0-N1-N2) and from panel ***B*** (HR + N0) are superimposed, red and blue, respectively. Of note, the peak shifts generated by N0-N1-N2 and N0 are well superimposed indicating that only OutD N0 domain interacts with OutB HR.

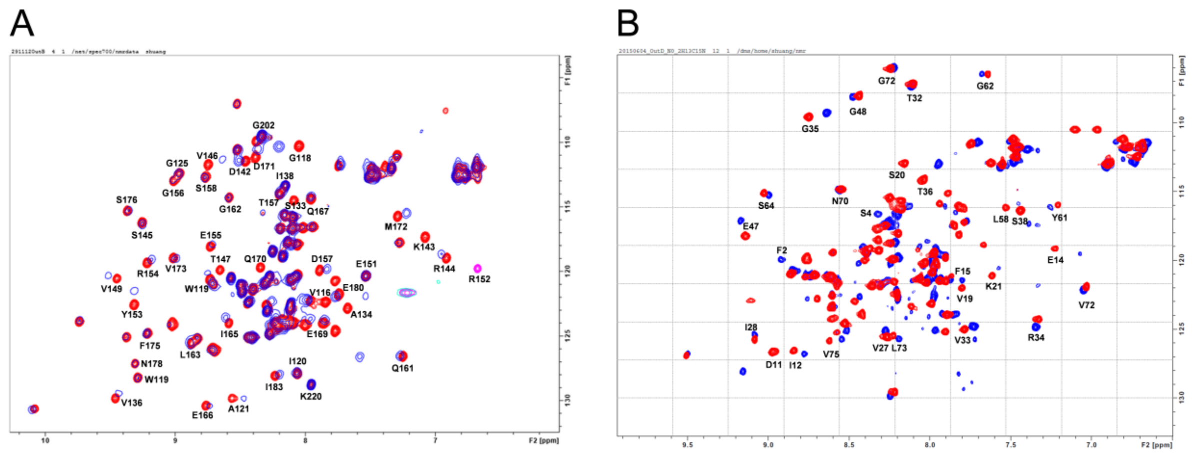


**Figure S5. NMR spectroscopy analysis of the OutB HR /OutD N0 complex. A.** $^{1}$H-$^{15}$N HSQC spectra of $^{15}$N-labelled OutB HR domain (residues P112-K220) alone (red) and with unlabeled OutD N0 (blue). **B.** $^{1}$H-$^{15}$N HSQC spectra of $^{15}$N-labelled OutD N0 domain (residues A1-S85) alone (blue) and with unlabeled OutB HR (red). Note that upon binding of HR, almost all $^{1}$H-$^{15}$N signals of N0 shifted illustrating that chemical perturbation method is not appropriate to determine the HR/N0 interface.

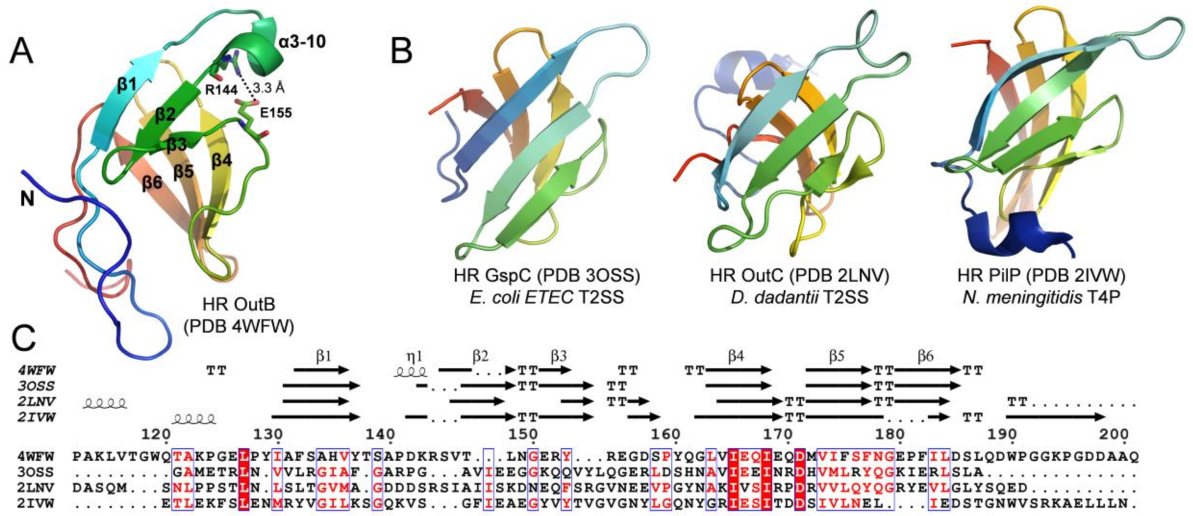


**Figure S6. The conserved periplasmic domain of OutB adopts the same topology as the HR domains of GspC T2SS and PilP T4P. A.** The crystal structure of the OutB HR domain (residues 112 to 202, PDB code 4WFW) is formed by two three-stranded anti-parallel β-sheets located at approximately 70° to each other that form together a β-sandwich. The loop between strands β1 and β2 forms a $3_{10}$-helix that is stabilized by a salt bridge between highly conserved Arg144 and Glu155. **B.** Some structural homologues of the *D. dadantii* OutB HR: *E. coli* ETEC GspC HR (PDB code 3OSS), *D. dadantii* OutC HR (PDB code 2LNV) and *N. meningitidis* PilP (PDB code 2IVW). The cartoons are coloured from the N-terminal end (blue) to the C-terminal end (red) of the polypeptide chain. **C.** Structure based alignment of the HR domains shown in *A* and *B* was performed with ESPript server (Robert and Gouet 2014). The residue numbering is this for *D. dadantii* OutB. The secondary structure elements are shown for the aligned HR structures. The groups of identical and similar residues are highlighted in red or are in red, respectively.

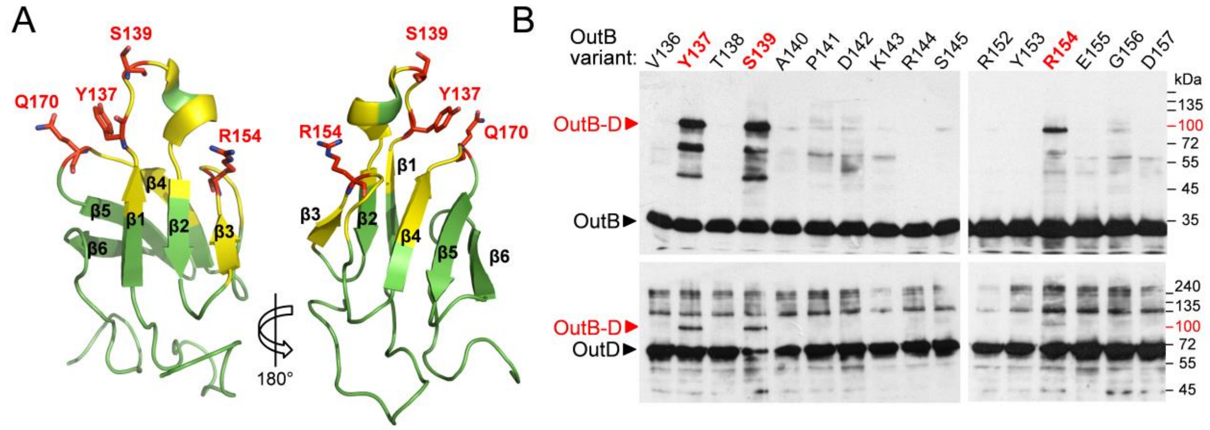


**Figure S7. Photo-crosslinking maps OutB residues interacting with OutD in *D. dadantii*. A.** OutB HR structure: the residues generating abundant OutB-D complexes are in red and other residues probed by photo-crosslinking are in yellow. **B**. *In vivo* site-specific photo-crosslinking. OutB *p*BPA substitutions (indicated at the top) were expressed from a DBS plasmid in *D. dadantii* Δ*outD* A3558/pREP4/pSupBPA. Cells were irradiated by UV (365 nm) for 3 min and analyzed by immunoblotting with anti-OutB and OutD antibodies, upper and lower panel, respectively. Only UV-irradiated cells extracts are shown. OutB, OutD and OutB-D complex are indicated by an arrow.

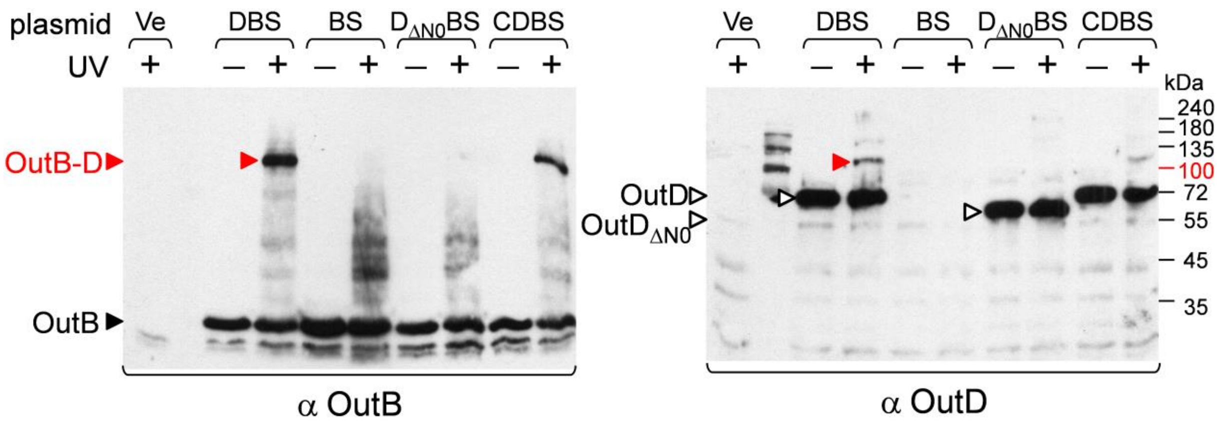


**Figure S8. Photo-crosslinking shows that OutB HR interacts directly with OutD N0**. OutB_R154*p*BPA were co-expressed together with OutS and with or without OutD, or OutDΔN0 and OutC in *D. dadantii* Δ*outD* A3558/pREP4/pSupBPA. Cells were irradiated by UV (365 nm) for 3 min and analyzed by immunoblotting with anti-OutB and OutD antibodies, left and right panels, respectively. The positions of OutB, OutD, OutB-D complexes and molecular mass standards are indicated.

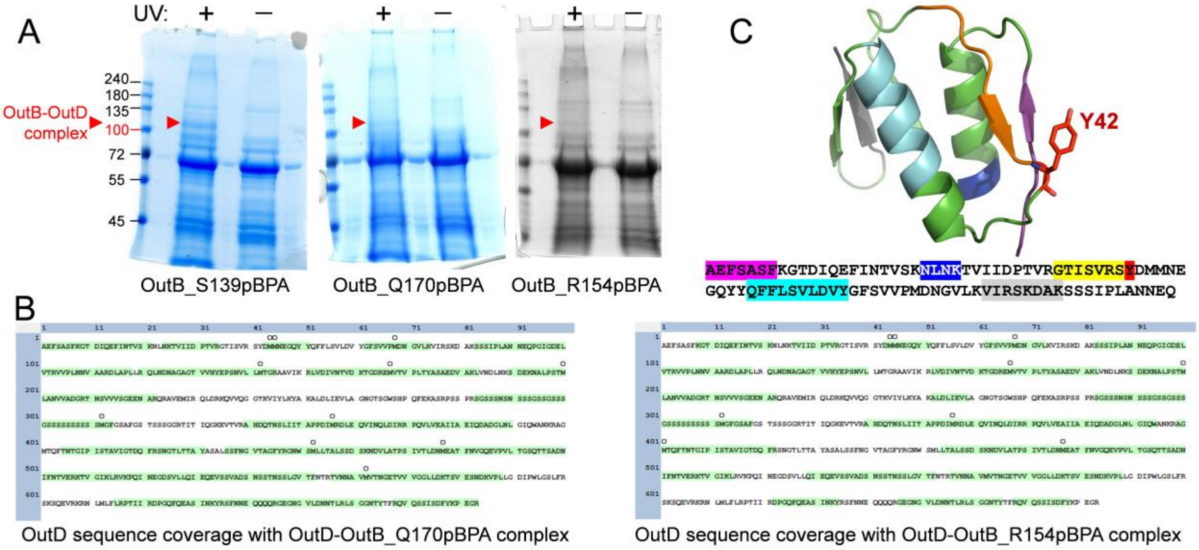


**Figure S9. Mass spectrum analysis of OutB_*p*BPA-OutD complexes.** **A.** *E. coli* cells co-expressing indicated OutB_*p*BPA variants with OutD-Strep were subjected to *in vivo* photo cross-linking and generated OutB-OutD complexes were purified by Strep-Tactin chromatography and separated by SDS-PAGE. The protein bands indicated with red triangles were excised and subjected to in-gel digestion with chymotrypsin and trypsin and analyzed by high resolution LC-MS/MS. **B.** OutD sequence coverage by the MS/MS analysis of the OutB-OutD complexes generated by OutB_Q170*p*BPA and OutB_R154*p*BPA variants. **C.** The peptides that were not assessed by MS/MS analysis in ***B*** are highlighted with different colors on the N0 OutD sequence and structure.

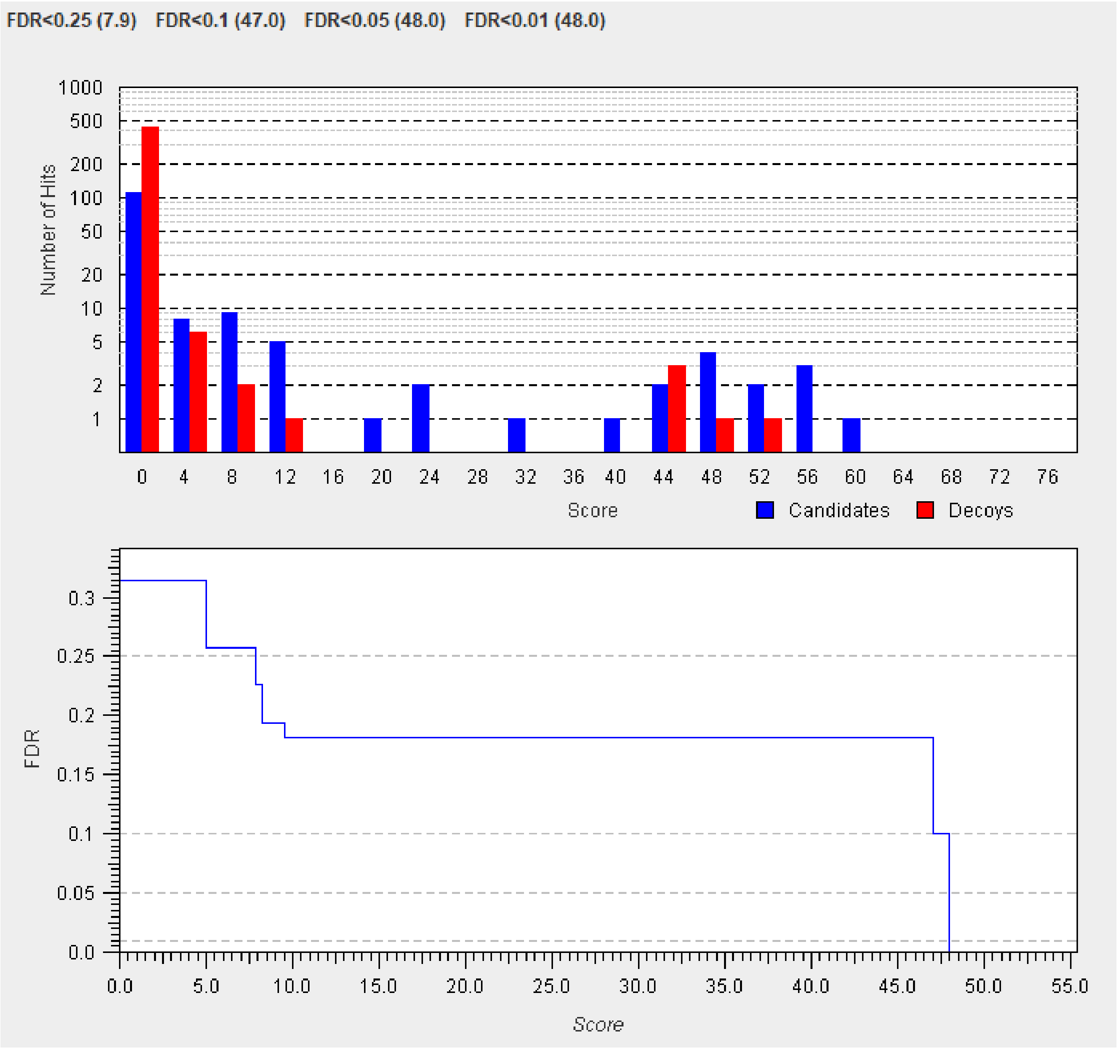


**Figure S10. StravoX validation of cross-linked peptide candidates by a Target-Decoy analysis.** The bar chart plots the number of candidates identified in a certain score range to the score range. The line graph shows the dependency of FDR at different score cut offs. A FDR of 1% is chosen corresponding to a score cut-off of 48.

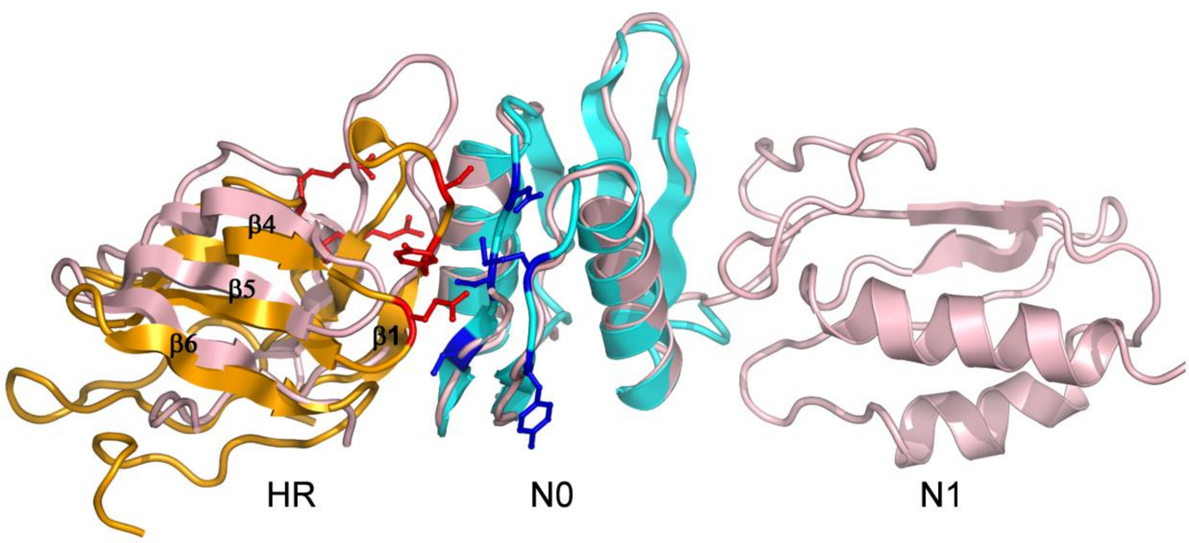


**Figure S11. Superimposition of the GspC HR/GspD N0-N1 complex (PDB entry 3OSS), in pink, with the OutB HR/OutD N0 complex, in gold and cyan, respectively**. The OutB HR and OutD N0 residues involved in the interdomain interface are shown in red and blue respectively. The superimposition is made by using only the N0 domains as templates.

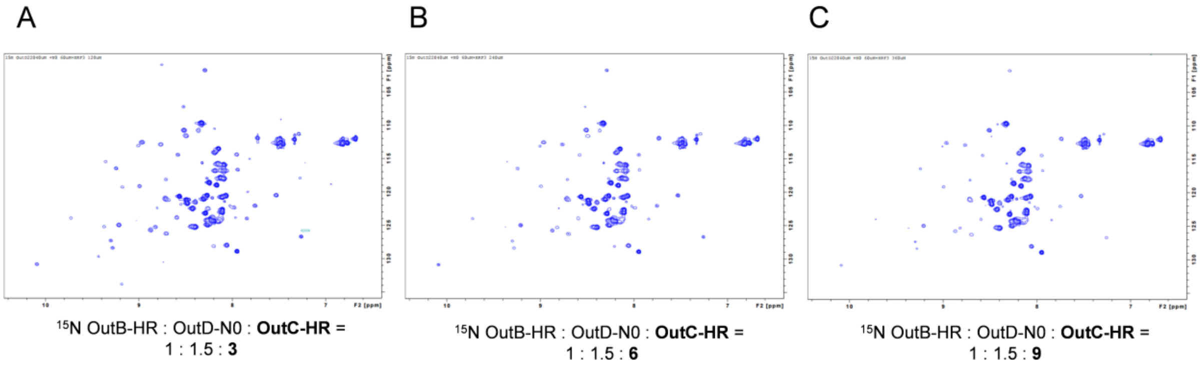


**Figure S12. OutC HR domain does not compete efficiently with OutB HR for OutD N0 domain *in vitro*.** Chemical shifts measured with $^{15}$N-labelled OutB HR domain (residues P112-K220) when unlabeled OutD N0 domain (residues A1-S85) and OutC HR domain (residues A78-S158) were added. The concentrations of proteins were as follow: $^{15}$N OutB HR, 40 µM; OutD N0, 60µM and OutC HR, 120, 240 and 360 µM, in panel A, B and C, respectively. No significant signal shifts were detected upon increased concentrations of OutC HR.

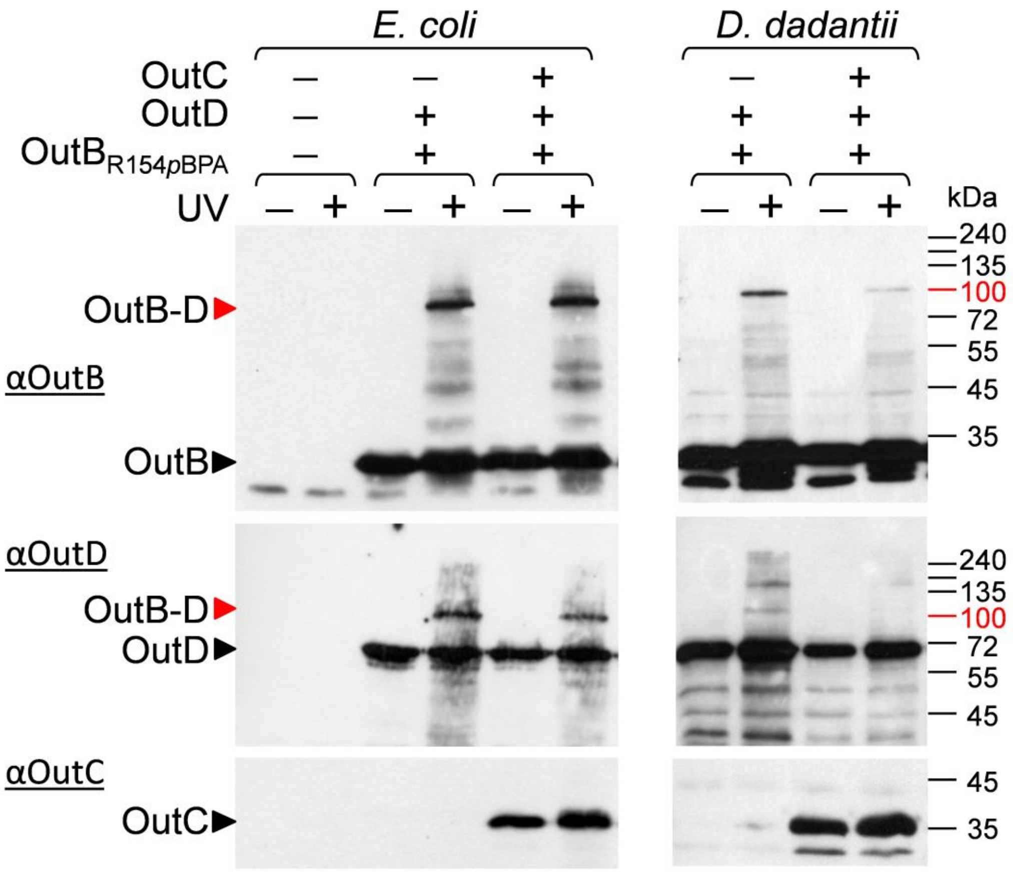


**Figure S13. In the functional T2SS, OutC and OutB could compete for the secretin OutD.** Effect of co-expression of OutC on the OutB-OutD interaction. OutB_R154*p*BPA variant was co-expressed with OutD and with or without OutC, either in *E. coli* MG1655/pREP4/pSUpBPA or *D. dadantii* A5654/pREP4/pSUpBPA, as indicated on top of the panels. Cells were irradiated by UV (365 nm) for 3 min and analyzed by immunoblotting with the antibodies against OutB, OutD and OutC. The data of a representative biological replicate are presented. In *D. dadantii*, the amount of OutB-OutD complex was steadily lower in the presence of OutC.

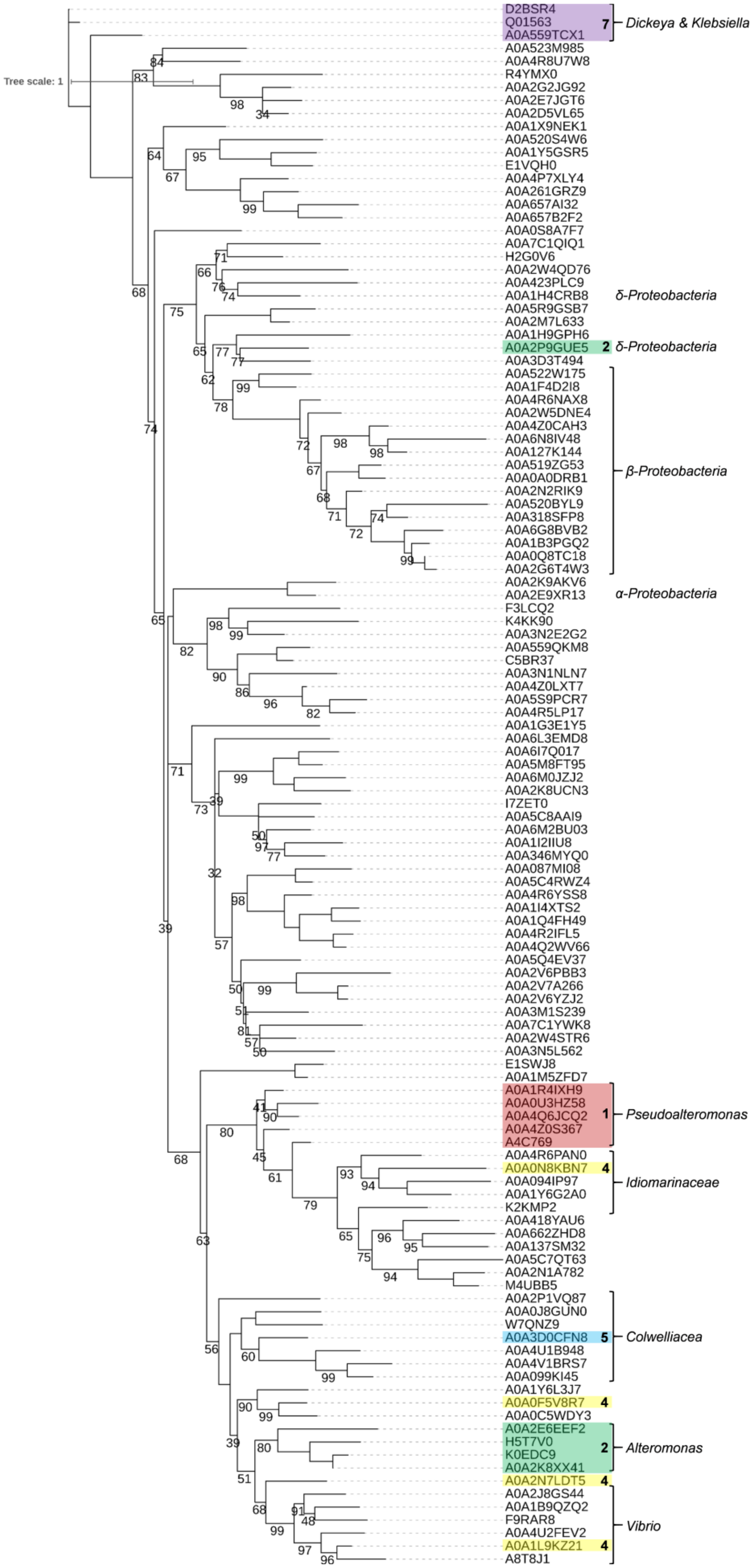
Tree scale: 1
D2BSR4
Q01563
A0A559TCX1
7
Dickeya & Klebsiella
A0A523M985
A0A4R8U7W8
R4YMX0
A0A2G2JG92
A0A2E7JGT6
A0A2D5VL65
A0A1X9NEK1
A0A520S4W6
A0A1Y5GSR5
E1VQH0
A0A4P7XLY4
A0A261GRZ9
A0A657AI32
A0A657B2F2
A0A0S8A7F7
A0A7C1QIQ1
H2G0V6
A0A2W4QD76
A0A423PLC9
A0A1H4CRB8
δ-Proteobacteria
A0A5R9GSB7
A0A2M7L633
A0A1H9GPH6
A0A2P9GUE5
2
δ-Proteobacteria
A0A3D3T494
A0A522W175
A0A1F4D2I8
A0A4R6NAX8
A0A2W5DNE4
A0A4Z0CAH3
A0A6N8IV48
A0A127K144
A0A519ZG53
A0A0A0DRB1
A0A2N2RIK9
A0A520BYL9
A0A318SFP8
A0A6G8BVB2
A0A1B3PGQ2
A0A0Q8TC18
A0A2G6T4W3
β-Proteobacteria
A0A2K9AKV6
A0A2E9XR13
α-Proteobacteria
F3LCQ2
K4KK90
A0A3N2E2G2
A0A559QKM8
C5BR37
A0A3N1NLN7
A0A4Z0LXT7
A0A5S9PCR7
A0A4R5LP17
A0A1G3E1Y5
A0A6L3EMD8
A0A6I7Q017
A0A5M8FT95
A0A6M0JZJ2
A0A2K8UCN3
I7ZET0
A0A5C8AAI9
A0A6M2BU03
A0A1I2IIU8
A0A346MYQ0
A0A087MI08
A0A5C4RWZ4
A0A4R6YSS8
A0A1I4XTS2
A0A1Q4FH49
A0A4R2IFL5
A0A4Q2WV66
A0A5Q4EV37
A0A2V6PBB3
A0A2V7A266
A0A2V6YZJ2
A0A3M1S239
A0A7C1YWK8
A0A2W4STR6
A0A3N5L562
E1SWJ8
A0A1M5ZFD7
A0A1R4IXH9
A0A0U3HZ58
A0A4Q6JCQ2
A0A4Z0S367
A4C769
1
Pseudoalteromonas
A0A4R6PAN0
A0A0N8KBN7
4
A0A094IP97
A0A1Y6G2A0
Idiomarinaceae
K2KMP2
A0A418YAU6
A0A662ZHD8
A0A137SM32
A0A5C7QT63
A0A2N1A782
M4UBB5
A0A2P1VQ87
A0A0J8GUN0
W7QNZ9
A0A3D0CFN8
5
Colwelliacea
A0A4U1B948
A0A4V1BRS7
A0A099KI45
A0A1Y6L3J7
A0A0F5V8R7
4
A0A0C5WDY3
A0A2E6EEF2
H5T7V0
K0EDC9
A0A2K8XX41
2
Alteromonas
A0A2N7LDT5
4
A0A2J8GS44
A0A1B9QZQ2
F9RAR8
A0A4U2FEV2
Vibrio
A0A1L9KZ21
4
A8T8J1

**Figure S14. Unrooted phylogeny of proteins carrying HR GspB domain (PF16537).** Phylogenetic tree topology was obtained by using bootstrapped maximum-likelihood approach with IQ tree web server as described in 'Experimental procedures' and visualized with iTOL software. Bootstrap values ≥35 are indicated below the branches. GspB proteins that represent archetypes #1, 2, 4, 5 and 7 (shown in Fig. 2B) are indicated and underlined with different colors. All the other, non-indicated cases represent the most abundant archetype #3. GspB proteins coming from α, β and δ-proteobacteria are indicated; the others, non-indicated, are from γ-proteobacteria. Note that GspA-B fusions (archetypes #4 and #5) appear in different groups.

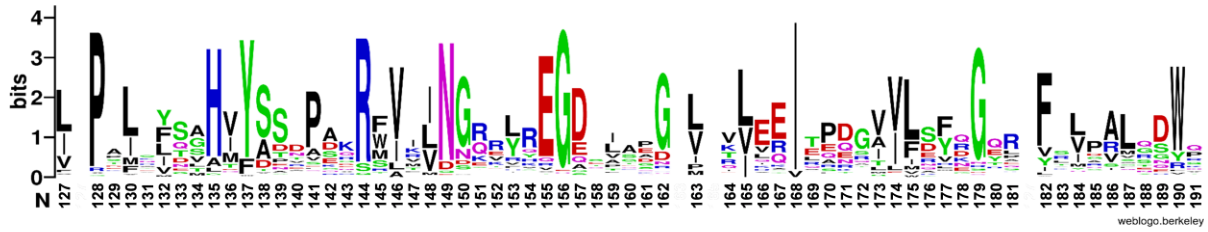

**Figure S15. Sequence logo of HR GspB domain sequences.** Graphical representation of multiple sequence alignment of HR GspB domains (117 sequences) used to generate the phylogenetic tree (Fig. S14). The height of symbols indicates the relative frequency of each amino acid at that position. Sequence logo was generated with weblogo.berkeley.edu server.

**Table S1. Data collection and refinement statistics**

| **Data collection** | |
|---|---|
| Space group | P $4_3$ $2_1$ 2 |
| Cell parameters (Å/°) | a=b=36.3, c=133.2 α=β=γ=90 |
| Molecules per asymmetric unit | 1 |
| Wavelength (Å) | 1.07 |
| Resolution (Å) | 2.02-36.30 (2.02-2.08) |
| Total number of observations | 56965 (2403) |
| Number of unique reflections | 6380 (435) |
| Multiplicity | 8.9 (5.5) |
| Completeness (%) | 99.2 (93.6) |
| $R_{merge}$ (%)[a] | 0.036 (0.574) |
| Mean <I/σ(I)> | 30.6 (3.7) |
| Wilson B-factor ($Å^2$) | 44.4 |
| $R_{pim}$(I) | 0.014 (0.280) |
| $R_{meas}$ | 0.041 (0.690) |
| **Refinement** | |
| Resolution limits (Å) | 2.05-34.91 |
| R-factor (%) / R-free[b] (%) | 20.85 / 25.42 |
| RMSD bonds (Å)/RMSD angle (°) | 0.015/1.81 |
| Average B-factor ($Å^2$) | 33.4 |
| Number of protein atoms | 666 |
| Number of solvent atoms | 15 |
| **Ramachandran plot statistics** | |
| Residues in most favoured regions (%) | 100 |
| Residues in additional allowed regions (%) | 0 |

The parameter values for the highest resolution shell are given in parentheses. The values presented in this Table are from SCALA (Evans 2006) and REFMAC from the CCP4 suite.

[a]$R_{merge} = \Sigma_{hkl} \Sigma_i \,|I_i - \langle I \rangle|\, / \Sigma_{hkl} \Sigma I_i$, where $I_i$ is the intensity of the $i^{th}$ observation, $\langle I \rangle$ is the mean intensity of the reflection and the summations extend over all unique reflections (hkl) and all equivalents (i), respectively. [b]R-factor = $\Sigma_{hkl} \,|F_o - F_c|\, / \Sigma_{hkl} F_o$, where $F_o$ and $F_c$ represent the observed and calculated structure factors, respectively. The R-Factor is calculated using 90% of the data included in refinement and R-free the 10% excluded.

**Table S2**. List of the best ten unique cross-links identified between OutB_S139pBPA and OutD_strep using StavroX analysis software.

| Score | m/z | Charge | M+H+ | Calculated Mass | Deviation in ppm | Peptide 1 | Protein 1 | From | To | Peptide2 | Protein 2 | From | To | Scan number | best linkage position peptide 1 | best linkage position peptide 2 |
|---|---|---|---|---|---|---|---|---|---|---|---|---|---|---|---|---|
| 63 | 780,00 | 3 | 2337,9821 | 2337,9791 | 1,31 | [SYDMMNEGQYY] | OutD-Strep | 41 | 51 | [TxAPDKR] | OutB_Strep_pBPA | 138 | 144 | 12591~190405-VS-0161-Chymo-Tryps-InclList01 | M4 | x2 |
| 59 | 780,00 | 3 | 2337,9811 | 2337,9791 | 0,84 | [SYDMMNEGQYY] | OutD-Strep | 41 | 51 | [TxAPDKR] | OutB_Strep_pBPA | 138 | 144 | 12143~190405-VS-0161-Chymo-Tryps-InclList01 | D3 | x2 |
| 57 | 780,00 | 3 | 2337,9818 | 2337,9791 | 1,15 | [SYDMMNEGQYY] | OutD-Strep | 41 | 51 | [TxAPDKR] | OutB_Strep_pBPA | 138 | 144 | 12704~190405-VS-0161-Chymo-Tryps-InclList01 | E7 | x2 |
| 56 | 780,00 | 3 | 2337,9792 | 2337,9791 | 0,05 | [SYDMMNEGQYY] | OutD-Strep | 41 | 51 | [TxAPDKR] | OutB_Strep_pBPA | 138 | 144 | 12816~190405-VS-0161-Chymo-Tryps-InclList01 | D3 | x2 |
| 54 | 544,48 | 4 | 2174,9169 | 2174,9158 | 0,52 | [SYDMMNEGQYY] | OutD-Strep | 41 | 50 | [TxAPDKR] | OutB_Strep_pBPA | 138 | 144 | 10690~190405-VS-0161-Chymo-Tryps-InclList01 | M4 | x2 |
| 52 | 585,25 | 4 | 2337,9806 | 2337,9791 | 0,65 | [SYDMMNEGQYY] | OutD-Strep | 41 | 51 | [TxAPDKR] | OutB_Strep_pBPA | 138 | 144 | 12594~190405-VS-0161-Chymo-Tryps-InclList01 | M4 | x2 |
| 51 | 585,25 | 4 | 2337,9804 | 2337,9791 | 0,55 | [SYDMMNEGQYY] | OutD-Strep | 41 | 51 | [TxAPDKR] | OutB_Strep_pBPA | 138 | 144 | 12159~190405-VS-0161-Chymo-Tryps-InclList01 | M4 | x2 |
| 50 | 725,64 | 3 | 2174,9172 | 2174,9158 | 0,66 | [SYDMMNEGQYY] | OutD-Strep | 41 | 50 | [TxAPDKR] | OutB_Strep_pBPA | 138 | 144 | 11407~190405-VS-0161-Chymo-Tryps-InclList01 | D3 | x2 |
| 49 | 725,64 | 3 | 2174,9176 | 2174,9158 | 0,83 | [SYDMMNEGQYY] | OutD-Strep | 41 | 50 | [TxAPDKR] | OutB_Strep_pBPA | 138 | 144 | 10767~190405-VS-0161-Chymo-Tryps-InclList01 | D3 | x2 |
| 49 | 780,00 | 3 | 2337,9811 | 2337,9791 | 0,84 | [SYDMMNEGQYY] | OutD-Strep | 41 | 51 | [TxAPDKR] | OutB_Strep_pBPA | 138 | 144 | 12255~190405-VS-0161-Chymo-Tryps-InclList01 | D3 | x2 |

**Table S3. Bacterial strains used in this study**

| Strain | Genotype/phenotype | Reference |
|---|---|---|
| *Escherichia coli* | | |
| BL21(DE3) | F$^-$ *dcm ompT hsdSB* ($r_B^-$, $m_B^-$) *gal lon λ* (DE3) | Stratagene |
| NM522 | *supE thi-1* Δ(*lac-proAB*) Δ(*mcrB-hsdSM*)*5* ($r_K^-$ $m_K^+$) | |
| | [F′ *proAB lacI*$^q$*Z*Δ*M15*] | NEB |
| MG1655 | K-12 F$^-$ λ$^-$ *ilvG*$^-$ *rfb-50 rph-1* | laboratory collection |
| MC4100 | F$^-$*araD139*Δ(*argF-lac*)*U169 deoC1 flbB5301 ptsF25 rbsR* | |
| | *relA1 rpsL150* | Casadaban, 1976 |
| MC3 | F$^-$*araD139*Δ(*argF-lac*)*U169 deoC1 flbB5301 ptsF25 rbsR* | |
| | *relA1 rpsL150* λ[ϕ(*pspA::lacZ*)] | Bergler *et al.*, 1994 |
| *Dickeya dadantii* 3937 | | |
| A3558 | *lacZ2 prt*$^-$ Δ*outD* | Bouley *et al.,* 2001 |
| A5652 | wild type | laboratory collection |
| A5653 | *outD::cat* ($Cm^R$) | This work |
| A5654 | *outB::uidA-nptI* ($Km^R$) | This work |
| A5719 | *outB::cat* ($Cm^R$) | This work |
| A5916 | outB::*nptI-sacB-sacR* ($Km^R$) | This work |
| A5954 | *outB*Δ*cte* | This work |
| A6078 | *outB::tetA* ($Tc^R$) | This work |

**Table S4. Plasmids used in this study**

| Plasmid | Genotype/phenotype | Reference |
|---|---|---|
| pGEM-T | P*lac*, P*T7pol*, *blaM* ($Ap^R$) | Promega |
| pET-20b(+) | P*T7pol*, coding PelB signal peptide and 6His, *blaM* ($Ap^R$) | Novagen |
| pSup-BpaRS-6TRN (pSup) | amber suppressor tRNA and aminoacyl-tRNA synthetase for incorporation of *p*BPA, *cat* ($Cm^R$) | Ryu & Schultz, 2006 |
| pREP4 | *lacI*$^q$, *neo* ($Km^R$) | Qiagen |
| pGEX-6P-3 | P*lac*, coding cleavable N-terminal GST, *blaM* ($Ap^R$) | GE (Helthcare) |
| CDBS [a] | pGM-T carrying *outC*, *outD*, *outB* and *outS* | This work |
| DBS [a] | pGM-T carrying *outD*, *outB* and *outS* | This work |
| DB [a] | pGM-T carrying *outD* and *outB* | This work |
| DS [a] | pGM-T carrying *outD* and *outS* | This work |
| D [a] | pGM-T carrying *outD* | This work |
| pGX-$oB_{HR\text{-}CTE}$ | pGEX-6P-3 carrying *GST-outB* (aa P112 to K220) | This work |
| pGX-$oB_{HR}$ | pGEX-6P-3 carrying *GST-outB* (aa P112 to G192) | This work |
| pGX-$oB_{HR2}$ | pGEX-6P-3 carrying *GST-outB* (aa P112 to G202) | This work |
| pGX-$oB_{HR\text{-}CTE}$-$oD_{N0\text{-}N1\text{-}N2}$ | pGEX-6P-3 carrying *GST-outB* (aa P112 to K220) followed by *N0-N1-N2-outD-6His* (aa A1 to V258) [b] | This work |
| pGX-$oB_{HR\text{-}CTE}$-$oD_{N0}$ | pGEX-6P-3 carrying *GST-outB* (aa P112 to K220) followed by *N0-outD-6His* (aa A1 to S85) [b] | This work |
| pGX-$oB_{HR\text{-}CTE}$-$oD_{N1\text{-}N2}$ | pGEX-6P-3 carrying *GST-outB* (aa P112 to K220) followed by *N1-N2-outD-6His* (aa A89 to V258) [b] | This work |
| pGX-$oB_{HR}$-$oD_{N0}$ | pGEX-6P-3 carrying *GST-outB* (aa P112 to G192) followed by *N0-outD-6His* (aa A1 to S85) [b] | This work |
| pET-oD28-112 | pET-20b carrying *N0-outD-6His* (aa A1 to S85) [b] | Login *et al.*, 2010 |
| pET-oD28-285 | pET-20b carrying *N0-N1-N2-outD-6His* (aa A1 to V258) [b] | Login *et al.*, 2010 |

[a] More details about these constructs are given on Fig. S2.

[b] Residue numbering is this for the matured, signal peptide-less OutD

**Table S5. Primers employed in the study**

| Primer | Nucleotide sequence (5'-3')[b] | Generated mutation[c] |
|---|---|---|
| OutB-BH-5' | **ctgggatcc**cccgccaaattggtaacag | |
| OutB-XB-RI-3' | **cggaattccggctctaga**gcatgatgtgcagttgctg | |
| OuB_H135TAG[a] | gtatatcgccttcagcgcg**t**a**g**gtctatacgtctgctccg | HR(OutB)H135tag |
| OuB_V136TAG[a] | cgccttcagcgcgcat**tag**tatacgtctgctccggac | HR(OutB)V136tag |
| OuB_Y137TAG[a] | ccttcagcgcgcatgtcta**g**acgtctgctccggacaag | HR(OutB)Y137tag |
| OuB_T138TAG[a] | cttcagcgcgcatgtctat**ta**gtctgctccggacaagcg | HR(OutB)T138tag |
| OuB_S139TAG[a] | gcgcgcatgtctatacgt**ag**gctccggacaagcgcagc | HR(OutB)S139tag |
| OuB_A140TAG[a] | gcgcatgtctatacgtct**tag**ccggacaagcgcagcgttac | HR(OutB)A140tag |
| OuB_P141TAG[a] | gcatgtctatacgtctgct**ta**ggacaagcgcagcgttacc | HR(OutB)P141tag |
| OuB_D142TAG[a] | gtctatacgtctgctccg**tag**aagcgcagcgttaccctg | HR(OutB)D142tag |
| OuB_K143TAG[a] | ctatacgtctgctccggac**t**agcgcagcgttaccctgaac | HR(OutB)K143tag |
| OuB_R144TAG[a] | cgtctgctccggacaag**tag**agcgttaccctgaacggag | HR(OutB)R144tag |
| OuB_S145TAG[a] | ctgctccggacaagcgc**tag**gttaccctgaacggagag | HR(OutB)S145tag |
| OuB_E151TAG[a] | gcgttaccctgaacgga**t**agcgctaccgtgaaggc | HR(OutB)E151tag |
| OuB_R152TAG[a] | gttaccctgaacggagag**tag**taccgtgaaggcgacagc | HR(OutB)R152tag |
| OuB_Y153TAG[a] | ctgaacggagagcgcta**g**cgtgaaggcgacagccc | HR(OutB)Y153tag |
| OuB_R154TAG[a] | ctgaacggagagcgctac**tag**gaaggcgacagcccgtatc | HR(OutB)R154tag |
| OuB_E155TAG[a] | gaacggagagcgctaccgt**ta**gggcgacagcccgtatcag | HR(OutB)E155tag |
| OuB_G156TAG[a] | ggagagcgctaccgtgaa**tag**gacagcccgtatcagggg | HR(OutB)G156tag |
| OuB_D157TAG[a] | gagcgctaccgtgaaggc**tag**agcccgtatcaggggttg | HR(OutB)D157tag |
| OuB_I165TAG[a] | ccgtatcaggggttggtg**tag**gagcagattgagcaggat | HR(OutB)I165tag |
| OuB_E166TAG[a] | gtatcaggggttggtgatc**t**agcagattgagcaggatatg | HR(OutB)E135tag |
| OuB_Q167TAG[a] | caggggttggtgatcgag**t**agattgagcaggatatgg | HR(OutB)Q135tag |
| OuB_I168TAG[a] | ggggttggtgatcgagcag**tag**gagcaggatatggtgatc | HR(OutB)I135tag |
| OuB_E169TAG[a] | gttggtgatcgagcagatt**t**agcaggatatggtgatcttc | HR(OutB)E135tag |
| OuB_Q170TAG[a] | gtgatcgagcagattgag**t**aggatatggtgatcttcag | HR(OutB)Q170tag |
| OuB_G193TGA[a] | gttgcaggattggccgggc**tga**aaaccgggcgatgacgcc | HR(OutB)G193tga |
| OuB_T214TAG[a] | cgtcaaaaccggagcaa**tag**gtcaggacaacgaagaaatg | CTE(OutB)T214tag |
| OuB_K219TAG[a] | caaaccgtcaggacaacg**t**agaaatgacacagcaac | CTE(OutB)K219tag |
| OuB_K219TGA[a] | gcaaaccgtcaggacaacg**tga**aaatgacacagcaactgcac | CTE(OutB)K219tga |
| OuB_Y137C[a] | ccttcagcgcgcatgtct**gc**acgtctgctccggacaagc | HR(OutB)Y137C |

| | | |
|---|---|---|
| OuB_T138C[a] | cttcagcgcgcatgtctat**tgc**tctgctccggacaagcgc | HR(OutB)T138C |
| OuB_S139C[a] | gcgcgcatgtctatacg*t**gc***gctccggacaagcgcagc | HR(OutB)S139C |
| OuB_Y153C[a] | cctgaacggagagcgct**g**ccgtgaaggcgacagcc | HR(OutB)Y153C |
| OuB_R154C[a] | ctgaacggagagcgctac**t**gtgaaggcgacagcccg | HR(OutB)R154C |
| OuD_E29TAG[a] | ccgtgtggctgggctgcc**t**a**g**ttttcagccagtttcaaagg | N0(OutD)E2tag |
| OuD_F30TAG[a] | gtggctgggctgccgaat**ag**tcagccagtttcaaaggaac | N0(OutD)F3tag |
| OuD_S31TAG[a] | ggctgggctgccgaatttt**ag**gccagtttcaaaggaacc | N0(OutD)S4tag |
| OuD_A32TAG[a] | ctgggctgccgaattttca**tag**agtttcaaaggaaccgata | N0(OutD)A5tag |
| OuD_S33TAG[a] | ggctgccgaattttcagcc**tag**ttcaaaggaaccgatattc | N0(OutD)S6tag |
| OuD_F34TAG[a] | gccgaattttcagccagtt**ag**aaaggaaccgatattcagg | N0(OutD)F7tag |
| OuD_R67TAG[a] | cgcggcaccatcagcgtg**tag**agttacgacatgatgaacg | N0(OutD)R40tag |
| OuD_S68TAG[a] | ggcaccatcagcgtgcgc**tag**tacgacatgatgaacgaag | N0(OutD)S41tag |
| OuD_Y69TAG[a] | ccatcagcgtgcgcagtta**g**gacatgatgaacgaaggg | N0(OutD)Y42tag |
| OuD_Y69C[a] | ccatcagcgtgcgcagtt**g**cgacatgatgaacgaaggg | N0(OutD)Y42C |
| OutDStrep | gccggcaacggaaccagcggc**tggagccacccgcagttcgaaaaa** | N3(OutD)WSHPQFEK |
| ROutDStrep | gggcgaggacggccgcgacgc**tttttcgaactgcgggtggctcca** | N3(OutD)WSHPQFEK |
| 5OuDdltN0 | **attttcagcc**ggtgatga gctggtgaccc | N0(OutD) ΔF7-I96 |
| 3OuDdltN0 | ctcatcaccggctgaaaa**ttcggcagcc** | N0(OutD) ΔF7-I96 |

[a] For each primers used in site directed mutagenesis, another primer with reverse complementary sequence was used (not shown).

[b] Mutated or introduced bases are in bold.

[c] Residue numbering is this for the matured, signal peptide-less OutD.

**Table S6. GspB representatives used in the sequence alignment on Fig. 2A**

| Short name [a] | Species | Accession number |
|---|---|---|
| OutB_Dd | *Dickeya dadantii* (strain 3937) | Q01563 |
| OutB_Ds | *Dickeya solani* | A0A2K8VW55 |
| OutB_Dz | *Dickeya zeae* | D2BSR4 |
| OutB_Da | *Dickeya aquatica* | A0A375A847 |
| OutB_Dc | *Dickeya paradisiaca* | WP_023638232 |
| GspB_Lb | *Lonsdalea britannica* | A0A1X3RVC9 |
| OutB_Pa | *Pectobacterium atrosepticum* | Q6D2I2 |
| OutB_Pc | *Pectobacterium carotovorum* | C6DAQ9 |
| PulB_Ko | *Klebsiella oxytoca* | A0A0H3H3J5 |
| PulB_Kp | *Klebsiella pneumoniae* | W9BIG8 |
| ExeB_As | *Aeromonas salmonicida* | A4SIF9 |
| ExeB_Ah | *Aeromonas hydrophila* | A0KPN3 |
| EpsB_Vp | *Vibrio parahaemolyticus* | Q87SK7 |
| EpsB_Vc | *Vibrio cholerae* | Q9KPC8 |
